# An integrated and scalable rodent cage system enabling continuous computer vision-based behavioral analysis and AI-enhanced digital biomarker development

**DOI:** 10.1101/2024.12.18.629281

**Authors:** Timothy L. Robertson, Michael Ellis, Natalie Bratcher-Petersen, Manuel E. Ruidiaz, Kevin Harada, Debra Toburen, Juan Pablo Oberhauser, Daniel Grzenda, Nicole E. Peltier, Madison Raza, Jan Benway, Jamie Kiros, Vivek Kumar

## Abstract

Home cage monitoring enables continuous observation of animals in familiar environments. It has large utility in preclinical testing, mechanistic studies, animal husbandry, and the general practice of the Replacement, Reduction, Refinement (3R) principles. Despite its acknowledged utility, home cage monitoring has not been broadly adopted. This is mainly due to the complexity of the tasks that must be solved to have a successful system that includes hardware and sensor development, data management, machine vision expertise, behavioral expertise, support, and user training. Here, we describe the Digital In Vivo System (DIV Sys), a modern end-to-end system for video-based rodent home cage monitoring. The DIV Sys consists of a cloud-based study design, monitoring, display, and visualization app (DIV App), local hardware for data acquisition cages (DAX), a machine learning model for tracking mice (mHydraNet) optimized for speed and accuracy, a study display and visualization app, and an advanced behavior quantification workbench (DIV Data). The platform seamlessly manages terabytes of video data in the cloud and is built around enterprise-level security and data standards. Collaborative tools enable teams across geographical locations to work together. As a demonstration of its utility, we used DIV Sys to analyze over a century of mouse videos across multiple geographic locations. We also characterized home cage behavior of 8 mouse strains and carried out customized video analysis. Together, we present a scalable home cage monitoring system for advanced behavior quantification for the rodent research community.

## 2 Background

Rodents, particularly mice, are the preferred species for modeling human diseases and conducting mechanistic and preclinical studies [1, 2]. While ideally, such studies would be carried out in natural habitats using ethological approaches [3, 4], modern laboratory experiments necessitate artificial housing environments and endophenotype-driven behavioral tests [5, 6]. However, current behavioral assays for modeling neurodevelopmental, neuropsychiatric, and neurological conditions face significant challenges in scalability, objectivity, reproducibility, and translatability [7–12]. The short duration and episodic nature of many behavioral assays often fail to capture the full spectrum of an animal’s behavior. These issues are well documented, with variability in results arising from differences in environment and procedures [7, 8, 13, 14]. These factors not only affect animal welfare, but also potentially skew mechanistic research outcomes and the translational relevance of animal studies.

In response to these challenges, researchers have increasingly turned to computational methods to study naturalistic behaviors as measures of health and disease [15–18]. Continuous capture and analysis of animal behavior data in a home environment, known as home cage monitoring, has emerged as a promising solution [19–23]. This approach offers several advantages, including assessment at the appropriate circadian phase, long-term observation potentially from birth to death, and minimal human interference [24, 25]. Observation of naturalistic, rather than elicited behaviors, is postulated to increase the reproducibility and generalizability of animal models [22, 23, 26, 27].

Animal behavior quantification has advanced rapidly with the application of machine learning (ML) and computer vision (CV) to the problem of behavior annotation and with the adoption of computational ethology [28, 29]. Often referred to as artificial intelligence (AI), these fields have benefited from breakthroughs in the fields of statistical learning and computer science, which have been adopted and extended for biological applications [30, 31]. Advanced methods have been used extensively in neurogenetic, preclinical, and numerous other studies. We have used computer vision methods to track visually diverse mice under complex environmental conditions, in gait analysis and posture analysis, to detect complex behaviors such as grooming, and to predict complex constructs such as frailty status, pain states, seizure severity, and animal mass [32–38]. Although these short-term studies have significantly contributed to our understanding of rodent behavior and genetics, there are limited applications of such high-resolution methods to long-term monitoring in home cages.

A wide range of technologies have been used to carry out home cage monitoring, including radio frequency identification (RFID), electrical capacitance, infrared beams, force plates, video, and hybrid approaches [20, 23, 39], and there have been multiple commercial and non-commercial systems developed for use with mice and rats. Each has advantages and disadvantages, and regardless of its acknowledged utility, wide adoption of home-cage monitoring has not occurred. One reason for that could be the complex nature of a successful platform [21, 22, 26]. A robust home cage monitoring platform necessitates several key components: specialized hardware and sensors, efficient data management systems, sophisticated machine learning models for behavior analysis, and accessible environments for data visualization and analysis. The establishment of such a hardware and digital ecosystem requires ongoing training, support, and maintenance. Consistent with the prevailing consensus in the field [26], commercially available systems are valuable for fostering broader utilization, enabling algorithm development, and providing sustained technical support. Given the advances in computer vision, video-based systems are the preferred modality to capture behavioral data and the sensor of choice for home cage monitoring. Hardware and enclosures must be readily available and meet the standards set forth in the Association for Assessment and Accreditation of Laboratory Animal Care (AAALAC) Guide for the Care and Use of Animals, while allowing high-quality data acquisition [40, 41]. The data load from 24/7 video monitoring is very large, with storage and management burdens. Data must be stored in an accessible environment for behavior analysis, since modern machine learning algorithms require large compute. Finally, algorithms for behavior quantification require advanced ML expertise. These models must be trained and validated across genetically diverse animals and varying environments, each with large visual diversity. All these components must be supported for the community over long periods of time. Thus, although seemingly easy, building a complex and scalable home cage monitoring system with proper hardware, software, data management, and algorithm infrastructure is a non-trivial task.

However, if properly executed, a digital monitoring platform has immense benefits through scalable and standardized studies, objective data analysis, and the ability to reuse data. Animal health and behavior can be uniformly monitored in an automated manner, providing greater sensitivity and likelihood of detecting signs of diease progression, therapeutic intervention, or impaired quality of life, thus improving animal welfare and likely study outcomes. From a neurogenetics and preclinical perspective, the models used to detect behavior, i.e. digital measures, can be developed and validated by and for the community. Key indices derived from multiple individual behaviors can be shared among users. The data are auditable, which is important in the preclinical space for ensuring research rigor and reproducibility, and supporting regulatory compliance (e.g. FDA). Finally, the ability to monitor animals continuously at high resolution to detect alterations in naturalistic behaviors has the potential to be more translationally relevant, thus improving drug development pipelines through improved predictive validity [16, 42].

Importantly, home cage monitoring aligns closely with the principles of the 3Rs in animal research: Replacement, Reduction, and Refinement [43]. Video data is reusable and multiple lines of research can be conducted with the same data, thereby enabling reduction in animal use. Improved sensitivity and specificity have the potential to lead to more insights gained from a single study, also leading to the potential for reduction in animal use overall. For instance, we used the same video dataset for tracking, gait/posture, grooming, and animal mass analysis [32, 33, 37, 38, 44]. Video-based monitoring of home cage behavior also leads to minimized handling requirements, thus reducing stress and for the observation of naturalistic behaviors not confounded by the presence of research personnel, or the white coat effect Adaptive study design and the comprehensive data collected can also potentially reduce the number of animals needed for studies, as more information can be obtained from each individual mouse [26, 45]. Continuous monitoring in the home cage combined with relevant digital measures can enable earlier detection of health and welfare issues, enabling more effective interventions.

With these factors in mind, we developed the Digital In Vivo System (DIV Sys). The DIV Sys is designed to address each of the components mentioned above in a modern cloud-based infrastructure. The platform has been developed by The Jackson Laboratory (JAX), a non-profit biomedical research institute with almost 100-year track record as a research and education resource for the biomedical community. The DIV Sys is designed to be a resource for the rodent research community that will be supported by JAX. It uses similar principles as the open field based JAX Animal Behavior System (JABS) [44] of data acquisition, data processing, and classifier sharing for behavior analysis with hardened hardware for data acquisition with data management and data analysis in the cloud, and study management, data visualization cross-platform in a web-based app in a home cage environment. It is a modular, turnkey system that does not require specialized knowledge to start while providing tools and resources for deep customized advanced behavior analysis. Six facilities across the U.S. and have been using it extensively for more than a year as a research application. In this report, we describe the system, present preliminary data, and present a path for community-driven adoption of advanced home cage monitoring. As these technologies evolve, they promise to revolutionize our approach to neuroscience, behavior, genetic, and preclinical research, to offer reliable, efficient, and ethically sound approaches for animal models.

## 3 Results

### Working Home Cage Definition

Although there are no strict definitions, Grieco et al. proposed preferable conditions for a home cage and home cage behaviors that include the ability to observe at least 1 light-dark (LD) cycle (although longer is preferable), and that the environment promote species-specific behaviors, has familiar bedding, food and drinking water, can have environmental enrichment, and includes cage ventilation for improved health status [21]. Others define it as the place where animals are housed for a majority of their lives [22]. The exact dimensions of the chamber can vary depending on the experimental objectives. These factors break down along a balance of space/facility considerations while promoting rich ethological behaviors. Systems have ranged from shoebox cages [46–49], to open field sizes such as JABS [21, 26, 44, 50, 51], to larger arenas [51, 52]. JABS for instance is can be used to monitor mice for weeks in a large open field environment. Larger dimensions require more space and enable a richer set of behaviors, while smaller arenas enable high-density husbandry with a limited behavior repertoire. Our requirements for cage size included meeting European Union space guidelines for the housing of mice, ability to use in high density rack systems, and the ability to repurpose for rats in the future [53]. We built the DIV Sys around a rat cage form factor that is a good compromise between our JABS field arenas and a shoebox mouse cage for rich behaviors. The system can collect data continuously for an indefinite period of time and has been used for several multi-month experiments.

### 3.1 Overall System Architecture (Figure 1A)

The Digital In Vivo System (DIV Sys) is an end-to-end system that includes an app for study design, hardware for video data acquisition, a machine learning pipeline, a data visualization portal, and an environment for advanced behavior analysis. These components fall into three systems components that are described in Figure 1. We briefly describe each in detail below and provide expanded details of our home cage monitoring system later in the results section.

**Figure 1:**
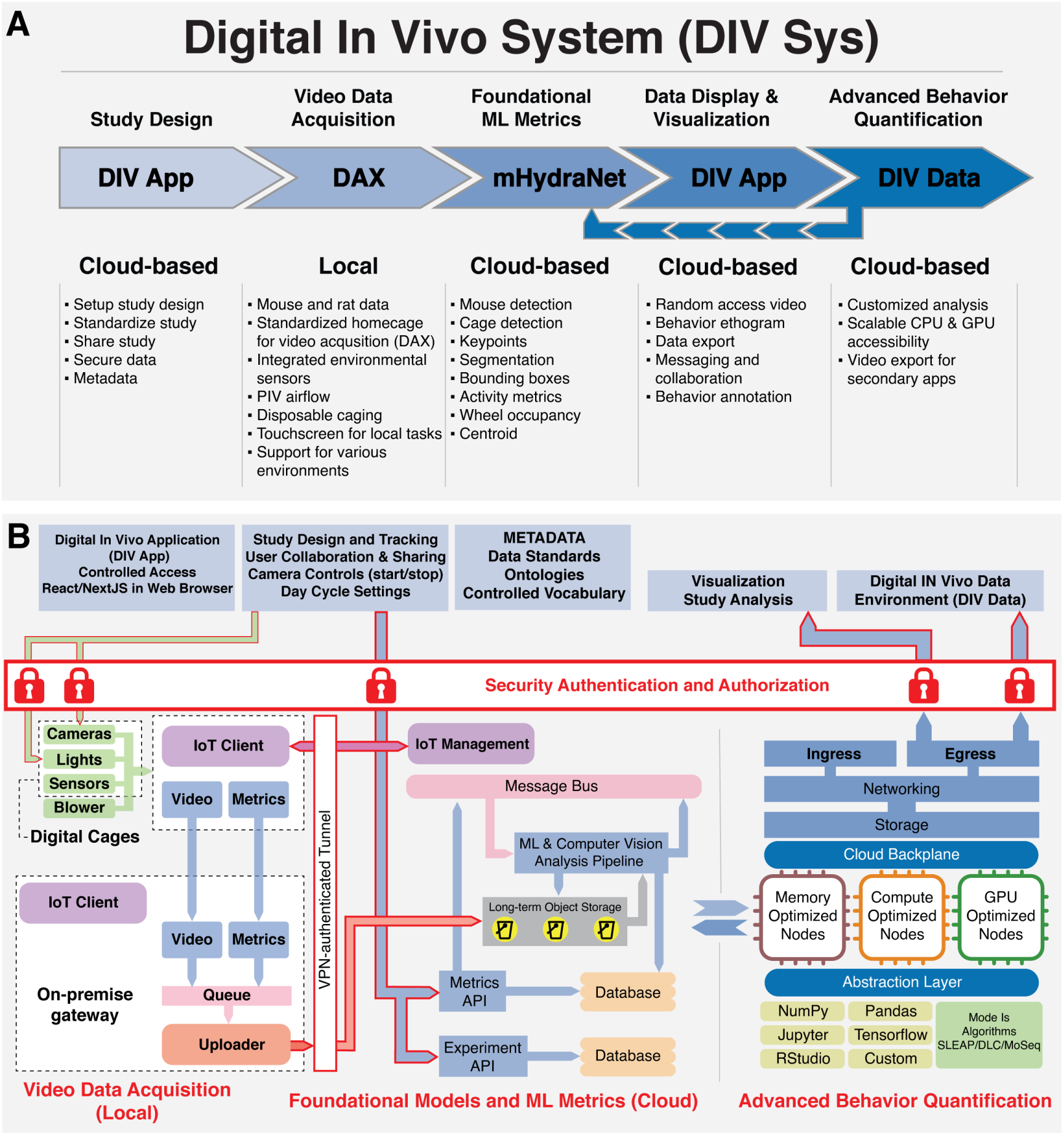
The Digital In Vivo System (DIV Sys), an end-to-end platform designed for behavioral research. **A.** DIV Sys Workflow Phases: A 5 step process including Study Design, Video Data Acquisition, Foundational ML Metrics, Data Display & Visualization and Advanced Behavior Quantification. *Study Design* A cloud-based interface to design and share experimental data securely. *Video Data Acquisition* Locally deployed hardware and software that captures high-quality video data and sensor data feeds from individual cages. *Foundational ML Metrics* Machine learning workflow leverages mHydraNet in the cloud to extract key behavior metrics. *Data Display & Visualization* An intuitive web interface allows detailed review and collaboration during and after data acquisition. *Advanced Behavior Quantification* The data environment to support advanced analysis for sophisticated behavior analysis with scalable cloud resources. **B.** System Architecture Implementation: Specific workflow functionality (top) enabled by DIV Sys Architecture in three principal segments: From left to right, these include: *Video Data Acquisition*, featuring on-site hardware such as sensors, networking gear, and digital enclosures; *Foundational Models and ML Metrics*, comprising cloud services and infrastructure for managing devices, storing data, and integrating APIs and databases; and *Advanced Behavior Quantification*, an analytical and developmental sphere for generating novel insights and analytics by researchers and data analysts.

#### Study Design (DIV App)

The Digital In Vivo Application is a cloud-based suite of capabilities that includes the design and management of digital studies (Figure 1A). The Digital In Vivo Application (DIV App) provides scientists and laboratory personnel with a secure browser-based interface to conduct and monitor their studies. Users can design studies, input metadata using a standardized vocabulary, and initiate recordings of their experimental cages. It is the first step in standardized and reproducible analysis. It enables standardized experiment design and captures essential metadata. Its functionalities allow users to annotate and share studies with collaborators in a secure data environment. It is fully described in Figure 2.

**Figure 2:**
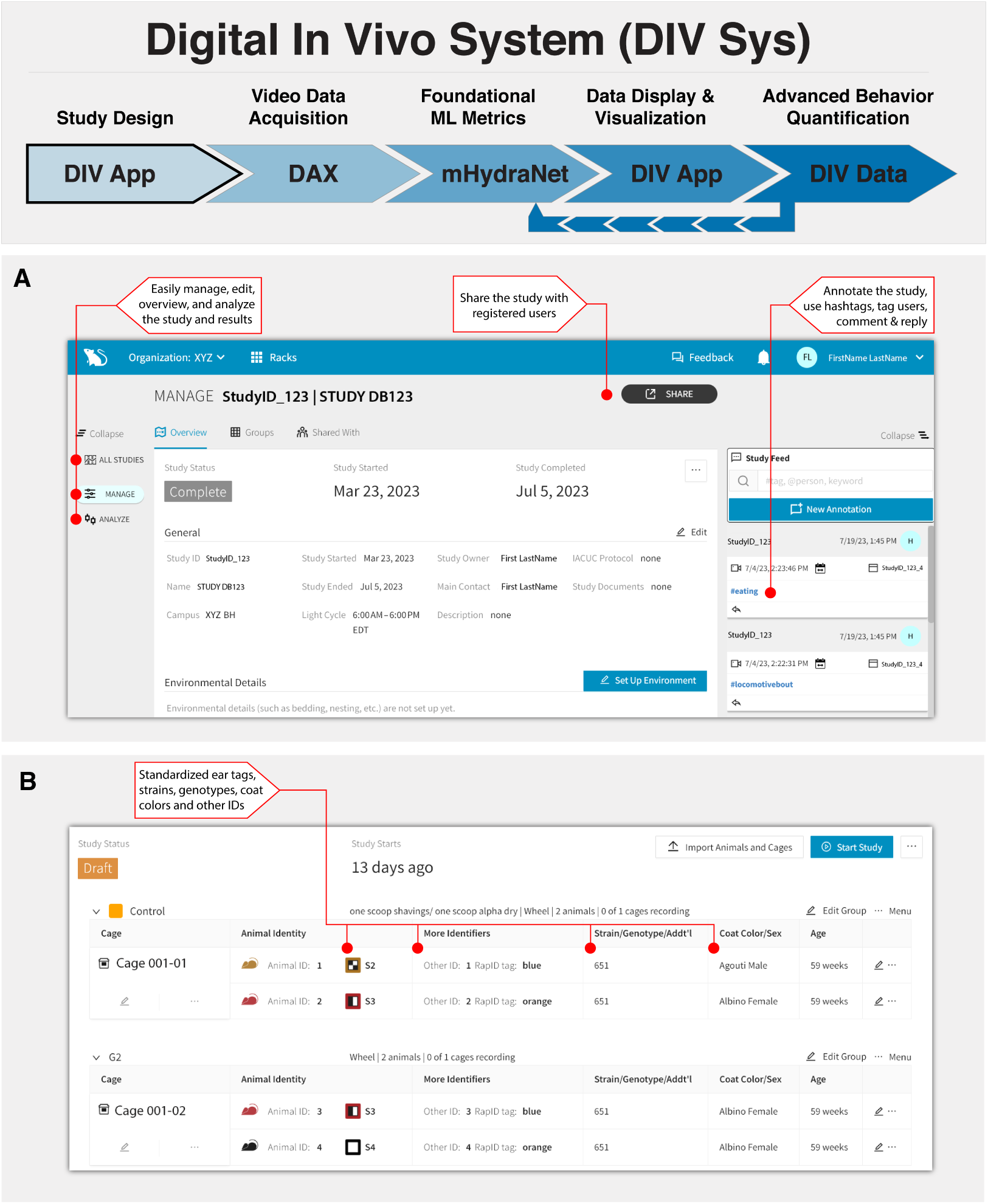
The Digital In Vivo Application (DIV App) **A.** Study design through the DIV App enables cloud based collaboration for standardized and documented studies. Study Design interface allows users to design and track studies. The application supports team collaboration with unique logins for each user. Scientists can design the full study using managed vocabulary to track the microenvironment (bedding and nesting used, enrichment, etc.) as well as animals. **B.** Animal metadata includes identification method, strain, genotype, age and other important details which may feed into other integrated systems. Technicians can execute the study and track progress, and can easily document activities and observations in annotations. Hashtags, mentioning other users, and replies support this collaboration. Managers can also access the system and quickly get an overview of all active studies, and ensure that the promised data is being generated. Scientists can share the study with other registered users, to support further collaboration.

#### Video Data Acquisition Hardware (DAX)

We have built a customized cage stall suitable for both mice and rats called the Digital Acquisition cage (DAX) (Figure 1A). DAXs are the primary data acquisition tool and can be deployed at geographically distinct facilities from experiment designers and analysts, disentangling these functionalities. The DAX housing enclosure has cameras, environmental sensors, and an ergonomic design that makes it suitable for vivarium integration. Standard disposable cages slide in and out of the stall without requiring any change to typical vivarium practice. These units are remotely managed and configured via cloud services to control environmental parameters such as lighting cycles and camera operations. The DAX unit is described in detail in Figure 3.

**Figure 3:**
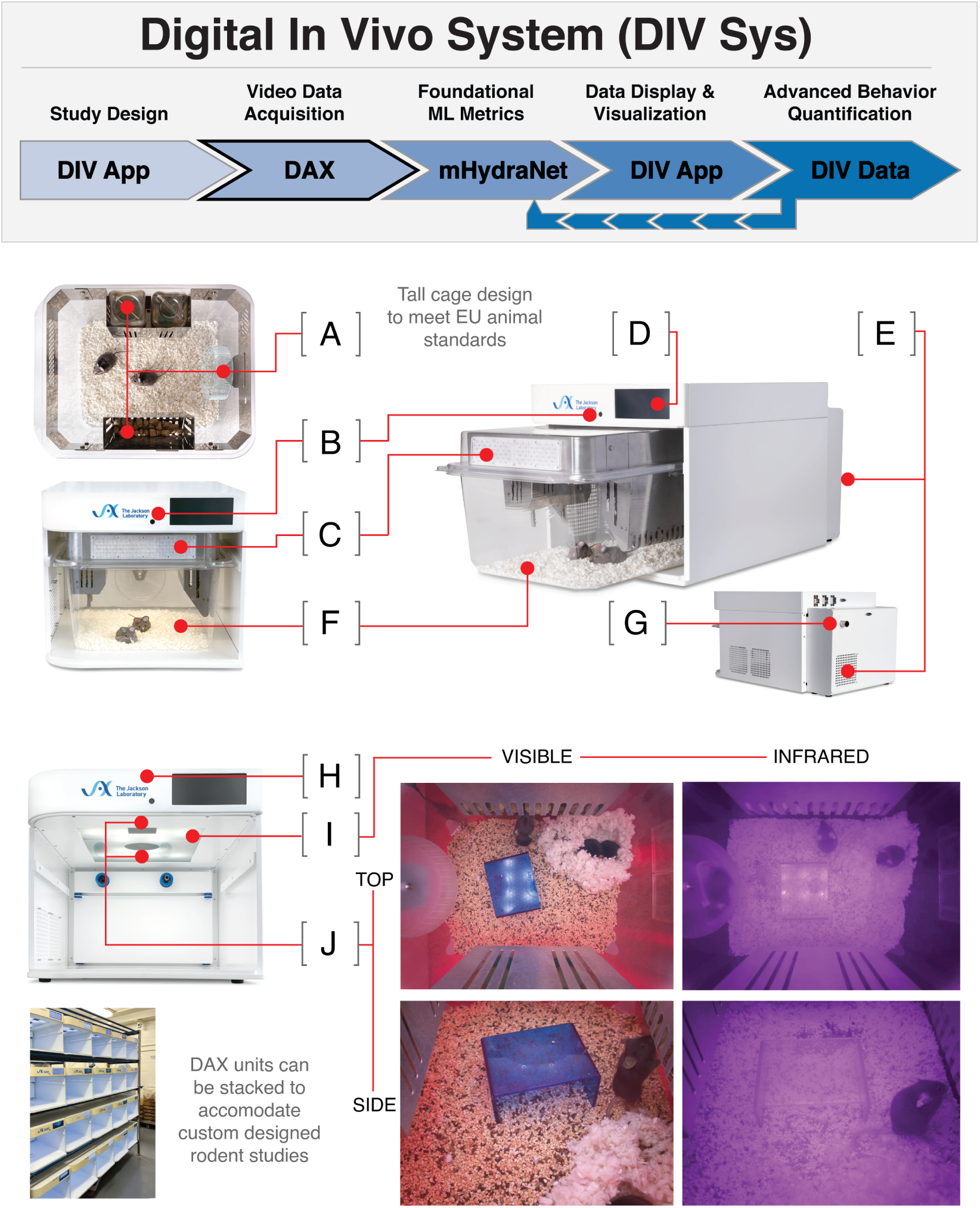
Video Data Acquisition through the DAX **A.** Integrated water bottles, food hopper, and running wheel, **B.** Light sensors to measure illumination, **C.** Vent for static ventilation if necessary, **D.** Touchscreen for digital cage card and basic data input, **E.** Vibration-isolated fans and HEPA filters in a backpack blower, **F.** Based on commercially available disposable rat bottom cages to support socially housed mice and rats, **G.** Data and power through a single POE cable, **H.** Jetson single-board computer for control, data acquisition, and local processing, **I.** Integrated visible and infrared lighting, **J.** Two high-resolution cameras for watching mice continuously. See Supplementary Videos 1-4 for sample videos of albino and black mice in day and night **Bottom Left.** High density racks of DAXs in a JAX animal room.

#### Foundational machine learning Metrics (mHydraNet)

Once video is captured by the DAX, it is deposited in cloud-based object storage, where a set of foundational metrics are extracted using machine learning and computer vision methods (Figure 1A). This includes the detection of cage change events, a mouse-optimized HydraNet (mHydraNet) that is used to extract salient keypoints, segmentation masks, and bounding boxes for each animal visible per frame. From this frame-level data stream, activity metrics are extracted. Zones for eating, drinking, and wheel running are defined and occupancy in these areas at the cage level is calculated. This module is key in converting images into meaningful biological insights. This process is described in Figure 4.

**Figure 4:**
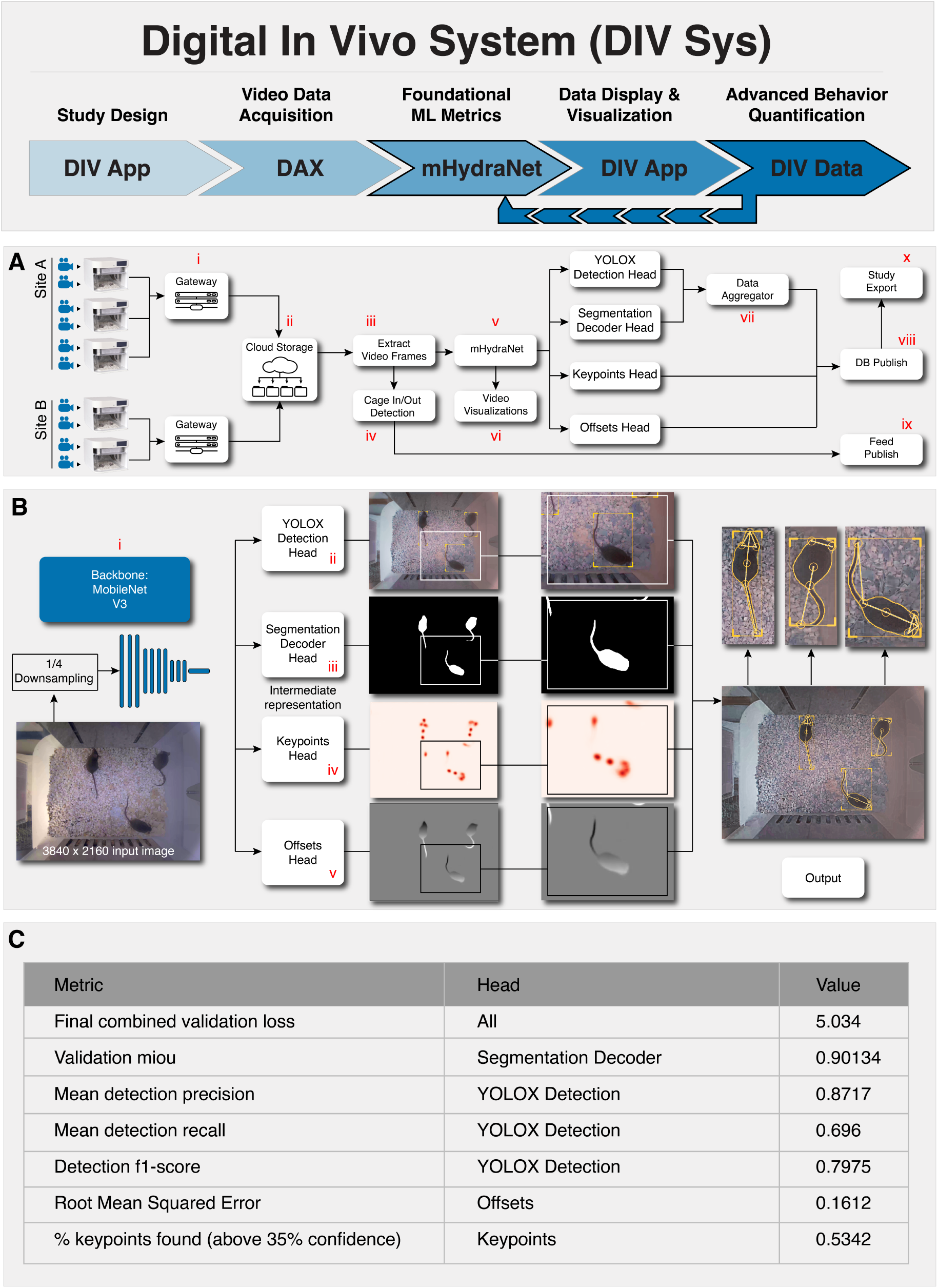
Data processing using Foundational ML Model (mHydraNet) for tracking of mice and environmental components. **A.** Data processing block diagram. Data from multiple sites feed into a uniform cloud storage and machine learning pipeline. The tracking data is aggregated and published in a study portal for viewing with DIV App. **B.** The main task of tracking mice is carried out by mHydraNet that extracts features from input images and outputs bounding boxes, segmentation, keypoints, and instancing offsets. The final output is shown on the right. **C.** Performance metrics of the mHydraNet.

#### Data Visualization Viewer (DIV App)

An accessible, intuitive and high-performing interface to view cloud-based video data is essential for behavior studies (Figure 1A). The foundational metrics are used to calculate behavior metrics that are displayed along with the primary video through a web-based interface. The application allows users to easily jump to and view any part of a video, facilitated by a modern playback engine. The system features customizable time-series plots which visually represent and track individual behaviors over time. Video overlays of mHydraNet outputs allow the user to inspect data underlying the various metrics traces. Data can be exported for further analysis. Messaging, collaboration, and annotation tools have been added to facilitate experimental and scientific collaboration. The data display and visualization app is described in detail in Figure 5.

**Figure 5:**
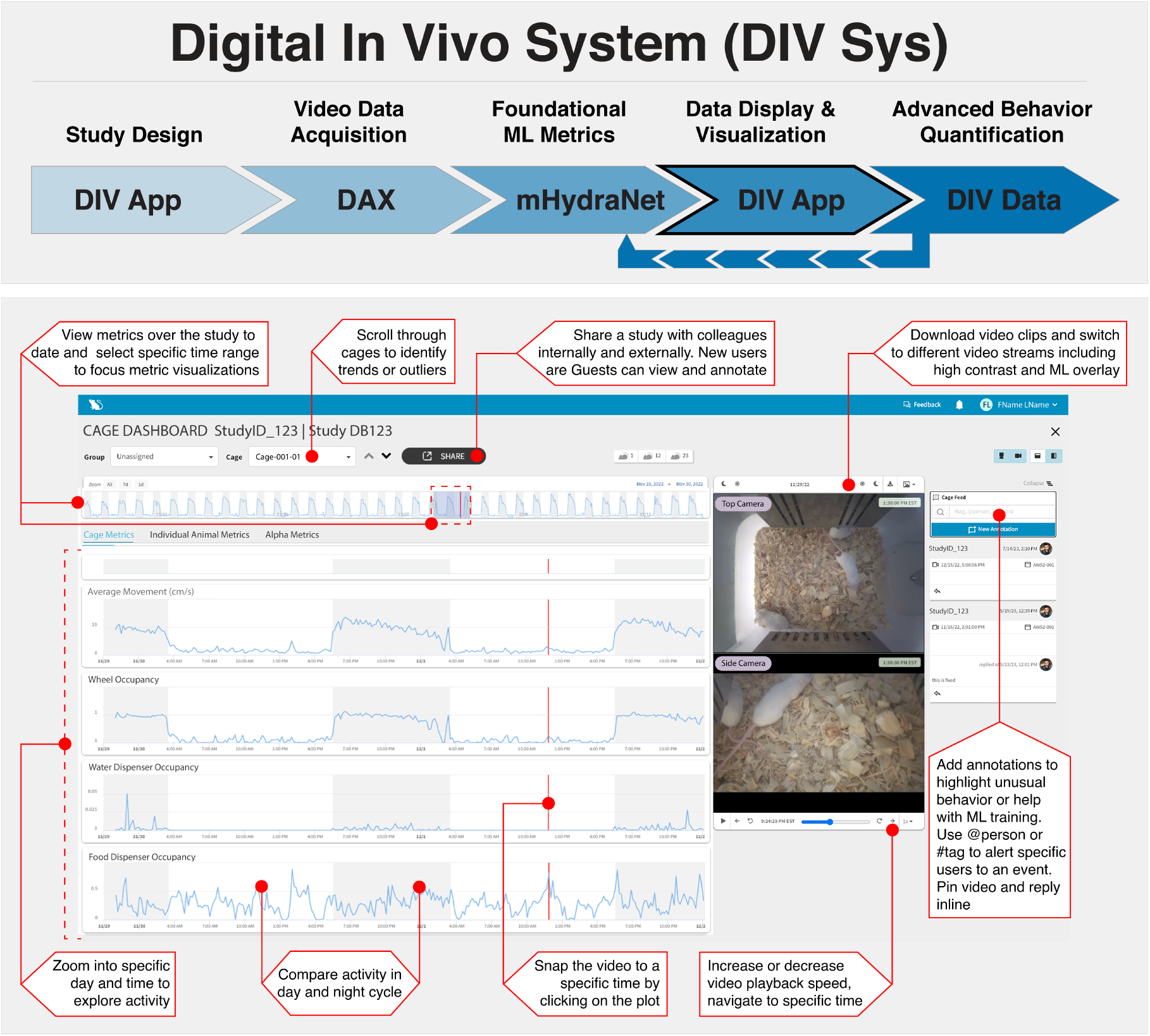
The DIV App enables video and behavior observation in an interactive and collaborative environment. Tools for behavior viewing, video playback and download, and tools for conducting and analysis collaborative studies. Over 30 days of data from a single cage are shown on the top ribbon. Bottom four ribbons show approximately 2 days of data for activity (average movement), wheel, water, and food dispenser occupancy. Windows on the right show top and side view videos. Annotations with @person or with `#tag` allows teams to communicate. HD video snippets can be downloaded or shared with collaborators.

#### Digital In Vivo Data Environment (DIV Data)

Another advantage of well curated data in the cloud is the ability to carry out advanced and customized behavior analysis. This is enabled by the Digital In Vivo Data Environment (DIV Data). This cloud-based tool provides scalable CPU and GPU access to carry out secondary analysis on the video data collected by the DAXs. It also links the DIV Sys platform with secondary algorithms that the field has and will continue to develop. Three applications of DIV Data are shown in Figures 6 and 7.

**Figure 6:**
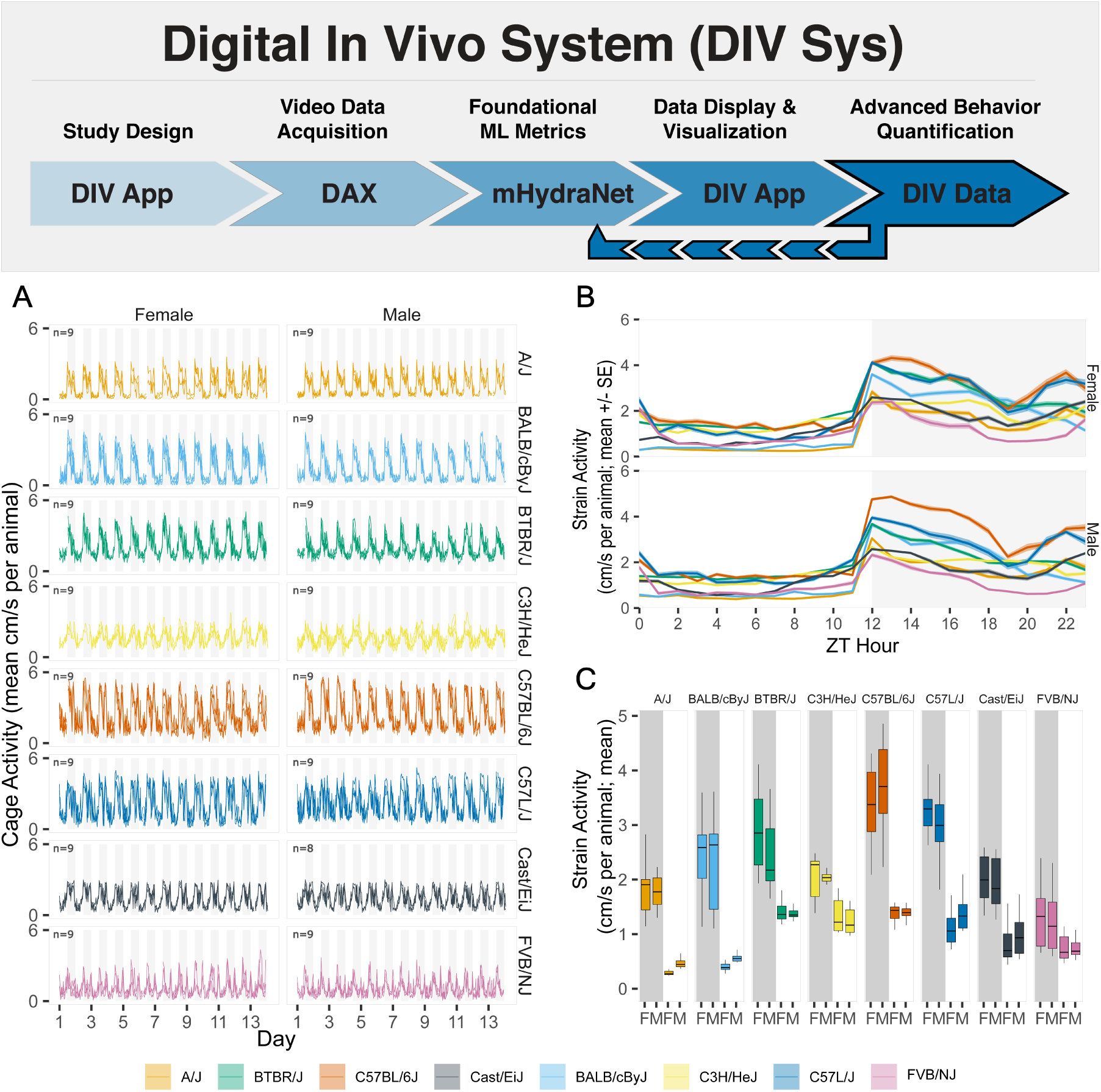
DIV Data allows custom advanced behavior analysis. In this vignette we characterize home cage activity of 8 mouse strains over a two-week period. (A). Individual mouse activity per cage is shown as movement speed (mean; cm/s) per hour over a 14 day recording period. Three groups of trio mice are shown for each sex and strain. (B) Diurnal variation in activity is shown as movement speed (cm/s; mean ±standard error) per hour derived from individual mouse activity over the recording period. (C) Average activity in night and day is shown as average speed and derived from the diurnal variation in activity. Boxes indicate quantiles for 12 hour interval.

**Figure 7:**
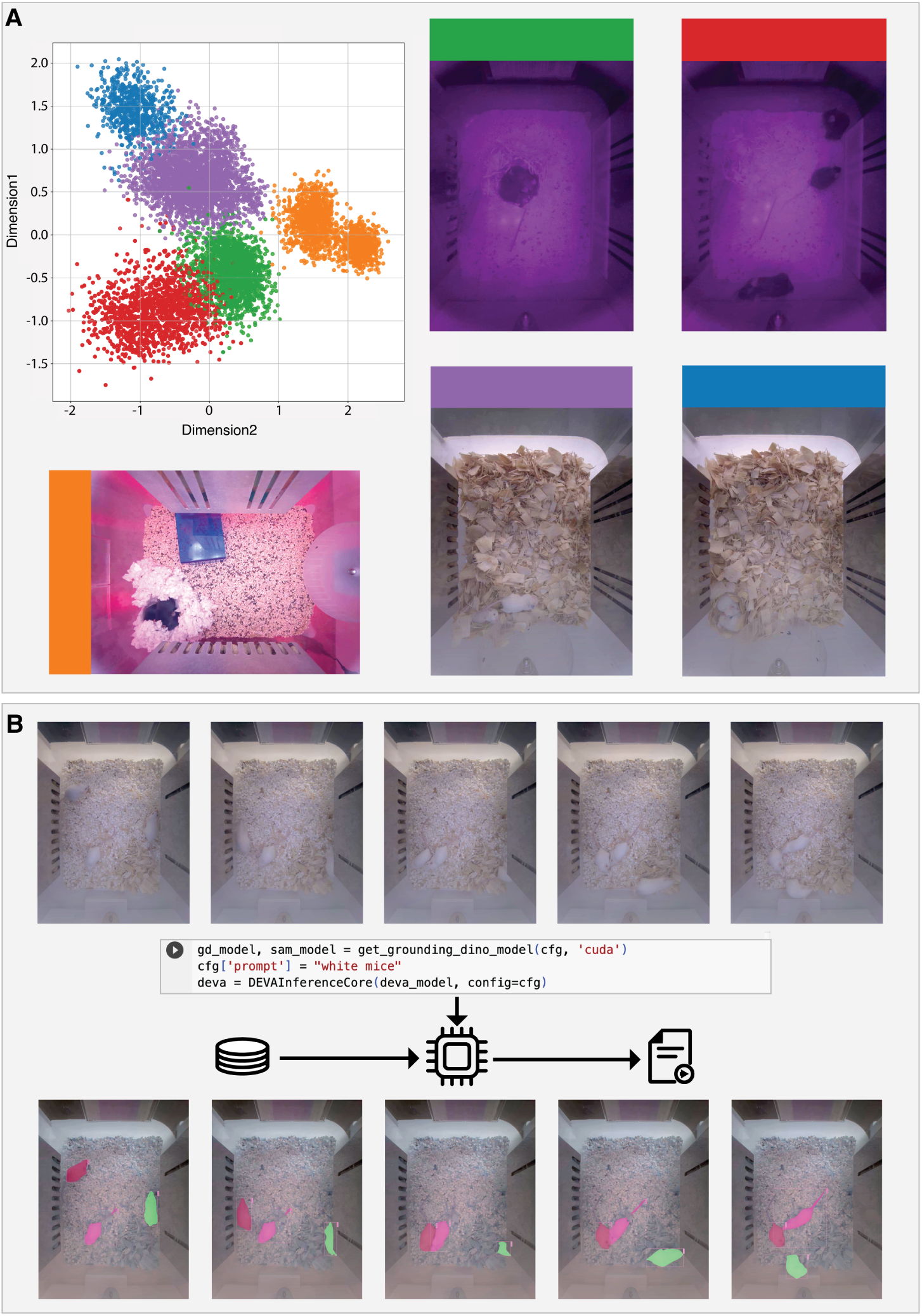
Advanced behavior and video analysis using DIV Data. **A.** Unsupervised analysis of 5 videos under varying states. Embeddings from mHydraNet of sample video frames are color coded and clustered. **B.** Zero shot segmentation using a SAM/DINO/DEVA pipeline to track individual mice (Supplementary Video 5)

### 3.2 System Architecture (Figure 1B)

The DIV Sys Digital Acquisition platform was architected by a group of software developers, machine learning practitioners, and domain experts with experience in cloud computing, web design, and internet of things (IoT) deployments, utilizing 12 factor app [54] design principles to ensure a robust software-as-a-service implementation. The technology stack is primarily implemented in Python [55] with a front end utilizing NextJS [56]. Containerization is extensively used to improve isolation and portability. The entire system is described using engineering block diagram style in Figure 1B.

The overall system can be divided into three major components, starting from left to right: 1) *Video Data Acquisition* includes on-premise hardware including physical sensors, remote networking equipment, and DAXs, 2) *Foundational Models and ML Metrics* includes cloud-based services and pipelines to enable device management processes, data storage, API endpoints and database engines, and 3) *Advanced Behavior Quantification* which is an analysis and development environment to understand and create new analytics by scientists and data practitioners. These components are described in detail below.

#### Video Data Acquisition

The components at the video data acquisition stage mainly comprise hardware and software that is running at remote sites. The two main hardware devices are the Digital Cage (DAX) and the On-premise Gateway. The DAX consists of cameras, lighting, environmental sensors coordinated by an NVIDIA Jetson Nano [57] system-on-module (SOM). The SOM has a quad-core ARM Cortex-A57 processor with a GPU based on NVIDIA’s Maxwell architecture [58], 4GB of RAM, 16GB of on-board storage and hardware accelerated video encoding. The Ubuntu 18.04-based [59] operating system hosts an Amazon Web Services (AWS) IoT Greengrass [60] Management client which allows remote device software deployments and configuration management. The GStreamer video capture pipeline [61] orchestrates video capture from the IMX467 camera utilizing the hardware video encoder to create H.264 encoded MP4 video files split into 1 minute segments. In parallel, the metrics service collects live measurements of system temperatures, light levels, resource utilization, and blower details.

These video and metrics data streams are pushed to the On-premise Gateway that acts as an uploader to downstream cloud resources through an secure VPN tunnel. A single gateway can handle streams from multiple cages, which also has a local buffering feature to minimize data loss during internet outages. Similar to the DAX, an IoT Management Client enables remote software deployments and configuration management.

#### Foundational Model and ML Metrics

In the cloud environment, video data is stored in an AWS S3 bucket [62] and metrics data are transferred via a Metrics API implemented in the Django Framework [63] which is backed by a PostgreSQL [64] database managed by AWS Relational Database Service [65]. These metrics are then utilized by the systems operation team for downstream alerting and visualization via a Grafana [66] dashboard.

Upon receiving each new 1-minute video chunk in the S3 bucket, an event is sent through an Apache Kafka [67] message bus, which starts a pipeline that initiates machine learning and computer vision based analysis. The pipeline outputs are stored as a combination of S3 and database outputs, which comprise the derived behavioral metrics. The transfer between S3, databases, and respective APIs are secured with implementations that enforce stringent authentication and authorization protocols, requiring that access is strictly controlled, segregated between accounts with different access permissions, and data security is maintained. A parallel service in this environment is the IoT Management layer, which is implemented using cloud-hosted AWS IoT Greengrass service.

#### Advanced Behavior Quantification

The Advanced Behavior Quantification module utilizes these metrics, offering data practitioners the tools to conduct in-depth analyses. To enable this workflow, a cloud-based deployment of the cluster management tool, Kubernetes [68] is hosted on Amazon Elastic Kubernetes Service (EKS) [69]. Within the Kubernetes environment, compute, network ingress, network egress, and file storage are abstracted from the underlying cloud implementation to enable a flexible computing environment. Elastic pools of virtual machines are configured for three types of compute tasks, memory intensive processing, compute intensive processing, and GPU intensive processing. Resources are organized in this fashion due to the diversity of compute requirements stemming from the interdisciplinary nature of behavior quantification, and to optimize cloud compute cost.

The primary user interface for this tooling is Kubeflow [70] which simplifies the deployment of ML and compute workflows on Kubernetes infrastructure. The Kubeflow interface includes support for computation notebook environments combined with containerized deployments, which enables a literate programming environment for the development of novel workflows [71]. Two widely used notebook environments, Jupyter Notebooks [72] and RStudio [73], are supported. Within these notebooks common data analysis libraries can be used, including, but not limited to, NumPy [74], Pandas [75], TensorFlow [76], PyTorch [77], R [78] and custom domain-specific algorithms such as SLEAP [79], DLC [80], and MoSeq [81].

### 3.3 DIV App: Study Design and Standardization

The DIV App is a cloud-based suite of capabilities that includes the design and management of digital studies. Each study is defined by a series of meta-data that outline the start and end of the study, the location, and links to critical documents such as the IACUC protocol. The overall study design is defined by the number and type of groups, such as treatment and control groups. Various groups can be defined, with any number of digital cages assigned to each group and up to three animals per cage (Figure 2A). Animal details such as strain, genotype, coat color, sex, and birth date are also captured. In addition, important to the study design is the identification method of each animal. The system is flexible to capture multiple methods of identification of animals such as RFID, tail tattoo, or ear notch (Figure 2B). We designed a set of custom RapID ear tags that are visible from the DAX camera (Figure S4). This key meta-data can be entered manually, or quickly uploaded from a spreadsheet. This study design drives the data organization and visualizations in the data viewer and analysis portions of the DIV App.

Another key element in the study design is the capture of environmental details, including bedding and nesting types, any supportive care, or furniture. The collection of meta-data for this study is controlled by a standardized dictionary to ensure consistent data entry. Future considerations include integration and expansion of the dictionary with known ontologies and the ability for an organization to create their own custom dictionary terms.

Once a study has been designed, it is automatically saved in draft mode and is available to others for review. As a cloud-based application, other users can review the study, input additional details, or edit the study design. A study can be shared with other researchers within the organization or external to the organization. This makes for a secure portal to not only design and manage studies, but also collaborate on that design and the resulting data collected. In addition, an Annotations Feed (Figure 2A) enables users of the system to comment and record significant events such as dosing, animal health, and behaviors of interest. An automatic annotation engine, the ML Bot, automatically posts detected cage removal and insertion events. The ability to pin video to the annotation and notify users with @mention or indicating a `#tag` event increases collaboration opportunities. For digital biomarker development workflows, there is also a confirm/reject process for ML Bot-generated annotations to involve scientists in verification of the metric. Scientists can review the output of the algorithm and correlate it with the behavior of the animal observed directly in the video.

Once a study has started, technicians can check the welfare of the animals through live stream video and metrics on the Cage Dashboard. Managers and researchers can review the progress of the study and initial metrics at both the cage and individual animal level. Visualization tools and the rich annotation feed from both manual and automatic annotations create a shared understanding of not only animal health and welfare, but also study progress and expected end points.

### 3.4 Hardware for Data Acquisition

We organized the animal habitat design into three groups of specifications, similar to the complementary JABS manuscript [44]. Since this is an end-to-end system, we do not expect laboratories to replicate the DAX. However, we provide the appropriate requirements for data replication. The first group of specifications describes the data acquisition that must stay constant for downstream ML based analysis. The second group describes components that can be modified as long as they produce data that adheres to the first group. The third group describes components that do not affect compatibility with our machine learning algorithms. While we distinguish between abstract requirements in group 1 and specific hardware in group 2 that meets those requirements, we recommend that users of our algorithms use our specific hardware in group 2 to ensure compatibility.

Group 1 specifications include the camera viewpoint, the minimum camera resolution and frame rate, the field of view and imaging distance, the image quality, and the general appearance of the habitat. Group 2 design elements are flexible but impact the compatibility and include camera model and lens. The lens focal length will impact the field of view, chromatic aberration, and capture performance. The focus procedure focuses each focal plane at a specific distance to ensure consistency between units. Group 3 design elements with no impact on compatibility include the height of the habitat walls to prevent mice from escaping or to meet IACUC standards, animal husbandry concerns, other sensors such as light and temperature, mounting hardware, ergonomic considerations for the technician, power and data connection hardware and management.

#### Unit encasing

The housing utilizes a disposable cage, designed to meet European animal housing standards and to facilitate social housing. The cage is positioned in a dedicated stall, which contains air fans, filters, illumination, cameras, sensors, and the Jetson single board computer for control, data acquisition, and optional local processing or data cloud transfer. The unit can be identified and controlled by a touchscreen that acts as digital cage card and enables basic data input and readout.

#### Illumination

Illumination is one the most important considerations of any computer vision-based system. Since mice are nocturnal, the majority of interesting behaviors occur in the dark phase. Similar to JABS we use two light sources visible and infrared, in the DAX [44]. Most cameras are more sensitive at shorter infrared wavelength illumination, which leads to better image quality. However, near-infrared (NIR) LED between 800 and 900nm have a red hue that is clearly visible and may alter animal behavior [82]. To determine the best wavelength to collect video data, we carried out an entrainment experiment using the circadian wheel running assay. The DAX light board has 16 NIR LEDs that can be adjusted in power and wave length (850nm, 890mn, 940nm, and white light). We asked whether C57BL/6J mice entrain to each wavelength at full power. The light board is shown in Figure S1A, with all emitters at full power (Figure S1B). The cell phone camera used to capture Figure S1B image shows a red/pink hue at 850 and 890nm, but not at 940nm. We placed eight light boards in one circadian light cabinet as shown in Figure S1C to ensure enough light at each wavelength. We then observed wheel running behavior of each mouse for seven days in the white-light schedule (LD), 850 nm, 890 nm, 940 nm and complete dark (DD) (behavior schedule in Figure S2A, B). We reasoned that if mice start free running, they are unable to entrain to that wavelength of light (Figure S2B, C). As an example, we show actograms of two animals that show clear entrainment in LD and 850nm, partial entrainment at 890nm, and free running at 940nm and DD. Summary statistics using LD or DD as control show that only 940nm is identical to DD conditions (Figure S2C,D,E). These data provided clear evidence for the use of 940nm NIR LED illumination over 840 nm or 890 nm for the DAX.

Informed by these data, the cage unit is furnished with fully controllable cage illumination system that provides 5000k White LED illumination with 100 lux at cage floor during the day, and 940nm NIR LED illumination for imaging in the dark. The system has LED persistence that ensures no illumination change occurs during soft reboots and is enabled with an illumination sensor to determine LED state. We confirmed that light conditions do not cause damage to the eye through evaluations by a trained pathologist at the Jackson Laboratory (Figure S3). Measurements of thickness of the photoreceptor layer and outer nuclear layer were evaluated in 20 mice and showed no significant variation (Figure S3A-B). There was no evidence of retinal atrophy or other degenerative changes. While several mice had eyes with developmental anomalies independent of lighting, no significant altered histopathology or morphology was observed.

#### Optical Sensors: Cameras and Illumination

There are 2 cameras placed in different positions - top center and top side - capable of recording at 4032x3040 pixel resolution at 30 fps. In addition, the unit is equipped an light sensor that reads ambient light in both white and IR channels. The DAX system collects 2 video streams (straight down and side oblique) at 4k and 1080p resolution. Video is captured in full color during the day and grayscale under IR light at night. The combined data rates are around 2-3 GB/hour. Sample videos for albino and black mice from top-center and top-side cameras are shown in Supplementary Videos 1-4. As of 2024, we have about 100 of these systems operating at 6 sites around the USA. To date (June 2024), we have captured almost a century of mouse video across all these cages, or approximately 100 billion frames.

#### Air Handling

The Jackson Laboratory was a pioneer in the development and adoption of microisolator or pressurized individually ventilated caging (PIV/IVC) [83, 84]. Such caging has been shown to protect animals from pathogens, as well as caretakers and scientists from animal allergens [85]. The DAX is equipped with fully controllable cage airflow, a system that provides dedicated supply and exhaust fans with flow rates up to 20 liters per minute (LPM) (Figure 3). It is also equipped with high-grade 99.999% HEPA filters that can create positive or negative cage pressure. The unit also contains vibration and noise isolation and medical-grade flow sensors (Figure 3E). A cage top filter provides sufficient air flow and the impact of a static environment was assessed in case of a power loss (Figure 3C). We monitored carbon dioxide (CO_2_), oxygen (O_2_), ammonia (NH_3_), temperature, and relative humidity within the DAX. Figure S5 shows these measures over the 8-hour assessment without power. Under these conditions CO_2_ and humidity rose steadily and O_2_ decreased over the course of the 8-hours. The veterinarian conducting the assessments noted that by 6-8 hours static, the mice were lethargic, possibly showing early signs of tachypnea while at rest. We conclude that animals are fine for several hours without power in DAX caging. The efficiency of the cage airflow component was also tested over 6 weeks with CO_2_, NH_3_, temperature, and relative humidity remaining consistent from week to week under normal operation (Figure S6). These results were stable over this course and indicate that the DAX is well suited for long-term studies.

#### Water, Food, and Enrichment

The cage is designed with in-cage food, water, and running wheel placement to minimize occlusion of subjects from the cameras (Figure 3A). Side viewing slots are sized and located to minimize climbing and enable food and water level by animal care technicians. The cage includes a food tray that is sized for two 500-gram rats with a wide feeding area to minimize aggression, and a water tray that can hold up to 600 mL of water with two drinking locations. A running wheel provides insights on animal activity, while other objects (gnawing blocks, hiding structures) offer an enriched environment.

### 3.5 Foundational ML Model Metrics

#### 3.5.1 ML and Computer Vision Analysis

DAX units are distributed at multiple sites within and across institutions, with a centralized data hub (Figure 4A). The multi-site on-premise gateways, buffer and transfer MP4 video files from each DAX to cloud storage in 1-minute intervals (Figure 4A.i & 4A.ii). This placement in cloud storage triggers an event on the message bus that starts the ML and Computer Vision Analysis pipeline, which then orchestrates the Foundational ML Metrics through a custom neural network (mHydraNet) as described in Figure 4A.

To ensure efficient processing on standard cloud infrastructure while safeguarding against the loss of processing events due to software, data, or hardware failures, our pipeline incorporates several resilience mechanisms. Initially, we adopted NATS JetStream [86], a software offering distributed persistence for message streaming. This setup is clustered, with storage nodes replicated and spread across three availability zones for enhanced reliability. For every component, a specific consumer is registered within the message stream, adhering to stringent retention policies and message acknowledgment protocols to guarantee comprehensive message processing. Furthermore, we employ sentinel files in the object storage system to establish a persistent record of the processing state, serving as a definitive source of truth for the system’s status. Sentinel files, named `metadata.json`, include details of component runtime, version, and other pertinent details to enable reproducibility of processing conditions and adjustment of parameters in future analyses.

##### Extract Video Frames Component (Figure 4A.iii)

Initially, MP4 video frames are extracted into about 56 (1 minute of frames) intermediate Zarr [87] files, each having a 32-frame downscaled (1008 x 760px) image tensors optimized for passing into our downstream mHydraNet model. These are temporarily stored in S3 and utilized by downstream components (Cage In/Out Detection, mHydraNet, and Video Visualizations are described below) that require access to pixel data. A strategy employing multiple batched files is adopted to enable the simultaneous multithreaded loading of data in parallel to active processing of loaded files. This approach ensures efficient parallel data handling and enhances the overall throughput of the system, while maximizing resource utilization.

##### Cage In/Out Detection Component (Figure 4A.iv)

This component determines cage presence in the DAX hardware by analyzing video frames. We extract one frame per second using a multi-threaded loader `zarr`. Several classical CV cage detection methods were tested (Figure S7). Detection is based on per-second texture computation in video frames. Full cages with bedding and furniture create more textural detail than empty or absent cages (Figure S7A). This detail is measured as gray-level cooccurrence matrix (GLCM) homogeneity, representing texture information in grayscale images [88]. Homogeneity indicates the similarity of intensity values in close regions [89]; higher values mean uniform textures, such as the empty bottom of the DAX, while lower values indicate the presence of texture, such as a cage. GLCM homogeneity is computed from the center of the video frame. Strided cutouts, sampling every 16th pixel, balance texture information and processing speed, producing 32 x 32 pixel cutouts from 512 x 512 pixel frames (Figure S7A). Over 2000 frames from six cage configurations (full, empty, and absent during day and night) were tested. Homogeneity histograms showed GLCM homogeneity distinguishes between cage conditions, with consistently higher values when the cage is absent at all times (Figure S7B). A homogeneity cutoff of 0.41 separates cage presence with an ROC curve area of 1.0. The cage detection results are compiled into a data frame with timestamps and metadata, stored as a Parquet file on S3 for analysis [90].

##### mHydraNet Component (Figure 4A.v)

This component begins by setting up the computational environment, including specifying the S3 bucket for model storage and determining the execution device based on CUDA availability. The model is loaded from a checkpoint snapshot in S3 including downloading of necessary TensorRT compiled models for efficient execution. We emphasize the use of specialized hardware acceleration to improve performance. The model processes video frames in batches as described above, it applies a series of transformations and utilizing the TensorRT compiled models for object detection (**YOLOX Detection Head**), segmentation (**Segmentation Decoder Head**), keypoint detection (**Keypoints Head**), and other tasks (**Offsets Head**). Each frame is analyzed to identify objects of interest, such as food, water, and wheels, with the outputs including segmentations, centroids, detections, and keypoints. Results from the model are saved both as compressed NumPy artifacts and as JSON objects, detailing the detected objects’ positions, sizes, and classes. This dual-format storage facilitates downstream analysis and integration with other components of the processing pipeline. Throughout the process, detailed logging and timing mechanisms are employed to monitor performance and ensure that the system operates efficiently. The use of asynchronous programming patterns and thread management techniques underscores this. We describe this component in-depth in section 3.5.2 below.

##### Video Visualizations (Figure 4A.vi)

Component: To visualize the outputs of mHydraNet, we overlay model predictions, including bounding boxes, keypoints, and segmentations, onto the original video frames. This process leverages the artifacts generated by the Extract Video Frames and mHydraNet components. Frames are sequentially extracted in batches from the intermediate Zarr output using a multithreaded loader to maintain the original sequence. Overlay artifacts corresponding to the model’s predictions are drawn onto these frames using OpenCV’s drawing functions. Subsequently, these annotated frames are encoded into an MP4 video file using an h264 video encoder. This MP4 file is annotated with a HLS (HTTP Live Streaming) playlist to facilitate adaptive streaming capabilities. The final step involves uploading the processed video and its metadata to S3, ensuring the availability of the results for further analysis or long-term storage.

##### Data Aggregator (Figure 4A.vii)

Component: The aggregation of individual frame outputs from mHydraNet into a time series output is essential for conducting study-level analyses. This component is tasked with performing data transformations and aggregations on model output payloads, employing Dask [91], a parallel computing library. Through Dask, a sequence of data transformations is applied, resulting in aggregated insights that encompass the positional data of animals within video frames. This aggregation is instrumental in a thorough analysis of their movements and interactions over time. A pivotal aspect of this process involves generating tracklets from per-frame detections to ascertain the distances traversed by the mice. These tracklets are formulated by utilizing temporal and spatial data, thereby coherently linking individual detections across successive frames.

##### DB Publish (Figure 4A.viii) and Feed Publish (Figure 4A.ix)

Components: These components play crucial roles as data sinks within our data processing pipeline, primarily focusing on handling metrics and event detections generated by upstream processing. Specifically, they deal with time series metrics derived from various detections, such as cage in/out states identified by the Cage In/Out component and aggregated data from mHydraNet outputs. DB Publish is tasked with integrating time series metrics into the metrics database. It leverages a Django data model abstraction to efficiently manage the bulk insertion of time series outputs while ensuring the integrity and consistency of the stored data. Conversely, the Feed Publish component is designed to handle real-time notifications based on specific event detections, such as changes in the cage in/out state. Upon detecting such an event, this component triggers a notification message to be sent to the Feed Publish system. This system then communicates with the Feed API through a GraphQL endpoint [92], ensuring that these events are promptly reflected in the annotation feature of DIV App. This mechanism ensures that users are kept up-to-date with real-time events and metrics, enhancing the interactivity and utility of the DIV app’s annotation feature. By employing these components, our pipeline not only efficiently processes and stores time series metrics but also enhances user engagement through real-time updates and annotations.

##### Study Export (Figure 4A.x)

Component: This step focuses on user-triggered exporting and structuring of study data for analytical examination by end users. Its role is essential for aggregating and scrutinizing data across varied timeframes, which is fundamental to extracting study-level insights. The user selects the study, via the DIV App interface, they wish to export. This triggers a series of queries to the experiment API to retrieve detailed study information, such as organizational identifiers and the relationships between cohorts and cages. This provides context to generated data aggregations consistent with the underlying study’s structure, critical for accurate data handling by downstream external analysis environments and tooling. An SQLAlchemy query is applied against the metrics database for relevant study timeseries data. Data aggregation is adapted based on user specified export criteria, either by cage or cohort. Thus, we compute summary statistics for the metrics within each group, transforming the data into a wide format that encapsulates all study metrics. Moreover, the dataset is enriched with additional metadata, including cage names and cohort animal counts, enhancing its interpretability. The integration of day cycle data, by calculating local times and categorizing them into light or dark cycles, based on experimental parameters, provides further insights into animal behavior within the study context. The final steps involve organizing the wide-format dataset chronologically and saving it in CSV format with columns named for clarity and emphasizing the resolution and measurement specifics of each metric.

#### 3.5.2 mHydraNet Model

Behavioral models in animals necessitate the systematic observation and quantification of specific activities that predict outcomes in more sophisticated patterns of behavior. This is best accomplished through initial abstraction of video data to track the animals [28]. The animal tracking field is a specialized subfield of human pose estimation that remains state of the art [93].

Essential to any tracking are foundational metrics, or intermediate representations, that serve as the building blocks for advanced behavioral analysis. These metrics encompass the location of the animal within its environment, the location of key body parts, and the descriptors of the overall animal morphology captured at any given moment. In addition, we need environmental information such as food hopper, water lixit, wheel, among others. We use three intermediate representations in our models - bounding boxes, segmentation, and keypoints. We have used segmentation and keypoints for tracking animals at high resolution in challenging conditions [32, 33]. We chose five keypoints as a starting point in our pose model to balance the richness of the representation and the ease of creating training data: nose, ears, middle of back, base of tail, and tip of tail. Although there is a strong emphasis on accuracy metrics in any ML model, for production environments such as the DIV Sys that produces millions of frames per day, speed and throughput is just as critical. We custom designed and trained a model to meet our needs.

Our model called mHydraNet (mouse HydraNet) accepts individual frames from a video stream and makes predictions about mouse detections, instance segmentations, keypoints, and other objects location such as running wheel, water tubes, and feeding area (Figure 4B). This processes a full minute of 1008 x 760 video at 30 fps in 30.2 seconds (∼ 60 fps) on a T4 GPU for inference, making it useful for potential real-time collection of multi-animal foundational metrics. mHydraNet is a hydra-style [94] model that uses a shared backbone as a feature extractor that outputs a pyramid of feature maps that are then fed into different heads responsible for different prediction tasks (detection, keypoints, segmentation). The entire mHydraNet implementation has only 5.52 million parameters, making it lightweight enough to possibly run on-device or on the edge. Due to the single-frame nature of this model, each incoming frame can be independently processed therefore, inference can be distributed over batches across different time steps or multiple camera sources.

##### Feature extractor

Input images are fed to a convolutional feature extractor following the mobilenetv3 architecture. This architecture is a convolutional neural network that is specifically tuned to run on small hardware. Therefore, it is a very small feature extractor that can be run both on the cloud or on the edge. Depthwise separable convolutions and linear bottlenecks are two of the breakthroughs presented in the MobileNet architecture family [95].

##### YOLOX Detection Head. (Figure 4B.ii)

The bounding box detection head is based on YOLOX which an anchor-free object detector [96]. The current mHydraNet is trained to detect mice of multiple coat colors in multiple environments, including while on running wheel, food hopper, and water lixit.

##### Segmentation head. (Figure 4B.iii)

The segmentation head outputs a feature map that is the same size as the original resolution image, with each pixel being labeled as mouse or no mouse. We then run a decoder as a top-down pathway from the highest level of the pyramid, merging and up-sampling feature maps at each step, up to and including level 3 of the pyramid. The final prediction is obtained by up-sampling the output of the original frame resolution.

##### Keypoint head. (Figure 4B.iv)

The keypoint head predicts 6 points of interest, left ear, right ear, tip of nose, base of tail, tip of tail, and animal centroid. Each keypoint is represented as a 2D isotropic Gaussian whose mean is the keypoint coordinate and variance is a tunable hyperparameter. The decoding step is similar to the segmentation head merging and up-sampling feature maps.

##### Offsets head. (Figure 4B.v)

A challenge then arises when doing multi-animal segmentation where each pixel with a mouse label needs to be individualized. One could, for example, assign each pixel in the segmentation map to the closest centroid (where the centroid comes from the keypoint head). Since mice are not convex objects, this kind of instance segmentation would often fail. For example, the tail of a mouse might be very close to the centroid of another mouse. Furthermore, the boundaries of mice touching each other would be linear. To overcome this, an offsets heads is added to the model, and it is tasked with computing per-pixel votes of which centroid the model believes the pixel is closest to, and acts as a bias term when computing the nearest centroid of a pixel.

#### 3.5.3 mHydraNet Training and Validation

##### Continual iterative learning paradigm

Each DAX can potentially acquire over 2.5 million image frames each 24-hour period. To effectively utilize this resource, we have integrated a continual learning pipeline. This is designed to selectively sample from the live data stream, employing a set of defined criteria to guide the training data selection process.

An example of this training selection criteria may include when the number of animals detected does not match the number of animals expected in the cage, where the mask does not meet certain complexity requirements, or where keypoints were missed or misplaced. In this case, while the model makes live predictions, the system creates a basin of images based on exceptions of the above training data criterion.

For example, the **Percentage of Keypoints Found** metrics reports when the keypoint head successfully identifies keypoints with above 35% confidence in 53.42% of cases. Keypoints are specific, predefined points on an object of interest, such as the extremities or joints of animals, which are crucial for understanding their posture, movements, and behaviors. The percentage of keypoints found reflects the model’s ability to detect these critical points accurately, which is essential for detailed behavioral analysis.

The image basin can fill up quite quickly given the number of frames, and selectively sampling from the basin for re-training becomes paramount. To do so, we utilize image embeddings to cluster similar images close to each other and use stratified sampling in an attempt to sample diverse miss cases in a representative manner. This approach not only streamlines the learning process but also enhances the overall efficiency and efficacy of the model’s training regime, critical for advancing research in our field.

##### Training

The mHydraNet model is extremely lightweight, therefore, training the model is relatively fast, requiring a single A100 GPU for 10-12 hours. During model training, each head outputs a scalar loss and the total loss is the weighted sum of all the individual tasks losses. The coefficients for the final loss are considered tunable hyperparameters.

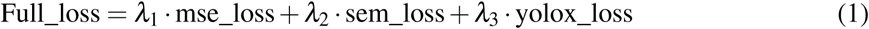

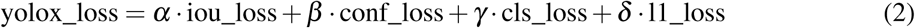

##### Continual training

We expect our model to continually evolve as more environments and organisms are sampled. The current reported results are of a model trained in late 2023. Reproducibility is very important for sharing or re-running certain studies in the system. Since the base model is being retrained on a cadence, the system keeps track of the version of the model that ran specific results, and every model can be re-used if necessary. The system also leverages a ‘run-id’ which keeps track of which model, which version of the code (tracked through github hashes), and which combination of components were run on a specific minute.

##### Final Combined Validation Loss

The final combined validation loss, with a value of 5.034, represents the aggregated error across all tasks the model performs. This metric is crucial, as it encapsulates the overall performance of the model, indicating how well it generalizes to unseen data. The loss is a weighted sum of individual losses from each model head, reflecting errors in segmentation, detection, and keypoint identification, among others. The weights for these losses are determined through hyperparameter tuning, aiming to balance the model’s focus across its various tasks. This approach ensures that the model does not overly prioritize one task at the expense of others.

Since we might not have data labels for every head, we use loss masking to enhance training data utilization. For example, if some data only have masks, we can still use it during training for keypoints if we flag it as missing keypoint data and mask the keypoint loss for those instances. The loss values for the final model are described in Figure 4C.

##### Validation YOLOX detection head

For the YOLOX Detection head, the Mean Detection Precision is 0.8717, and the Mean Detection Recall is 0.696, leading to an F1-score of 0.7975. Precision measures the accuracy of the predicted positive observations, while recall (sensitivity) assesses how well the model identifies all relevant instances.

##### Validation segmentation decoder

The Validation mean intersection over union (mIoU) for the Segmentation Decoder stands at 0.90134, indicating a high degree of accuracy in segmenting the target from the background. Intersection over Union (IoU) is a common metric for evaluating the accuracy of an object detector on a particular dataset. It measures the overlap between the predicted segmentation mask and the ground truth mask, with a higher score denoting better segmentation performance.

##### Root Mean Squared Error for Offsets

The Root Mean Squared Error (RMSE) for Offsets is reported at 0.1612. This metric quantifies the average magnitude of the error in predicting the offsets for detected objects. Offsets are vector distances from a detected object to a specific point of interest, crucial in tasks requiring precise localization within an image, such as tracking animal movements or identifying specific body parts.

##### Inference

Once the model is trained, the inference step has several sub-tasks to perform, including linking keypoints to individual animals. The first step is using the YOLOX detection head to output a collection of bounding box proposals for animals of interest. At the same time, the segmentation head outputs a proposal of animal vs no-animal per pixel. To obtain instance segmentation (different instance of animal), the model uses the centroids and offsets (where each pixel essentially votes for which centroid it is closest to with its offset vector). Lastly, for each detected animal, all keypoints are associated to the instance by conditioning them to the instance segmentation and box that have already been calculated. For inference acceleration, we leverage NVIDIAs TensorRT framework [97].

##### Derived Metrics

The outputs of the mHydra model are then used to begin to extract derived metrics that matter or give insight into specific behaviors. For example, average distance traveled, time on wheel, time near feeder, and time near water are metrics reported in real time per cage. The movement by animals in a cage in a given period of time is defined simply as the amount of distance that all animals travel in a given period of time, thereafter normalized and shown as a speed. This process runs a simple multi-object tracker that keeps track of individual tracklets and links bounding boxes and polygons across time. Then, the centroid of the polygon is calculated at each time step and the euclidean distance of the centroid movement is tracked across time. With a simple pixel to centimeter conversion, we get a measure of how many cm a tracklet moved across a period of time. Those tracklet distances are all added up to represent the cage-level distance traveled.

At the beginning of the study, a boundary box is recorded that signals the location of the running wheel, the feeding tubes, and the water tubes. Using that location, the first version of the time on "x" metrics is a measure of how much time a single animal bounding box centroid is within a bounding box of either the wheel, the feeding area, or the water tubes.

### 3.6 Digital In Vivo Application (DIV App) for Cloud based Data Visualization

The Digital In Vivo Application (DIV App) contains a graphical viewing portal for digital endpoints and video (Figure 5). We envision the DIV App as a central portal for metadata, behavior metrics, and a collaborative platform for researchers. The Cage Dashboard enables interrogation of the study data and video for the selected cage and animals from study start to current time or end of the study. The top navigation plot displays the activity metrics that have been collected for the life of the study and enables the user to select a subset of data, weeks, days, or minutes, to view on the tabs and four metric plots that are displayed below. There are drop-down menus to scroll through groups and each cage to compare activity or screen for animal health issues. The Dashboard includes multiple tabs under the top navigation plot that represent various digital end points. The first tab is the cage level metrics, the second tab represents individual animal metrics, and the third tab displays additional end points that are in the development phase enabling users to verify early metrics with the Confirm/Reject Annotation workflow. On each plot there is a red line indicating the placement of the video play head. Users can zoom in and pan on the individual plots and click a point to place the play head for the corresponding top and side camera to play. The gray shaded area represents the night cycle which is set specific to each organization. Metrics are aggregated depending on the zoom level the user selects. The example data in Figure 5 shows 36 days of data of continuous recording, with a detailed view of 48 hours of data.

The video player contains additional controls to navigate the video directly. Options include the ability to play forward or backward, snap to previous or next day or night cycle, and set the playback speed slower or faster enabling users to quickly scan for significant behaviors or look for animal health issues. The menu drop down contains multiple video display streams including a contrast enhanced video with animal ID overlay (when using the special ear tags) and high-resolution. Clicking on the video will bring that camera (top or side) to full screen mode to view more details. In addition, video clips can be downloaded at any time intervals.

On the far right of the Cage Dashboard is the Annotations feed which enables users to create comments and pin a specific time point in a video. In addition, users can be mentioned with an @mention which will send them an email and an in application notification alerting them to the mention. Links in the notification email will bring them directly to that point in the video where the annotation was made, making collaboration on specific studies and events easier. Annotations are also displayed on the Event Bar above the metrics plots. The Event Bar includes all annotations or can be filtered to display a specific subset enabling the correlation of animal activity metrics to annotation events such as cage removal/insertion, or dosing events. Clicking on the interactive annotation on the Event Bar snaps the video and annotation feed to that specific time point for quick review. The reply capability in the annotation feed enables direct communication and closer collaboration on events and projects.

### 3.7 Digital In Vivo Data Environment (DIV Data) for advanced behavior analysis

The final component of DIV Sys is a scalable platform for advanced behavior quantification called DIV Data. Video data are rich, enabling multiple levels of scientific interrogation; however, no one standardized analysis pipeline will meet the needs of all laboratories. The DIV Data platform allows users to access DAX data programmatically on a scalable cloud platform with CPU and GPU access and gives users virtually unlimited ability to carry out secondary and tertiary analysis. We provide three vignettes in which we used DIV Data to answer specific questions for advanced behavior analysis.

#### 3.7.1 Strain survey for home cage behavior

We used DIV Sys to characterize the home cage behavior of eight inbred strains of mice over a 2-week period. We selected strains of varying coat colors and sizes for analysis. Groups of three males and three females were placed in the DAX with a 12 hour light-dark cycle and with *ad libitum* food and water (see Table 1 in Methods). Individual mouse activity levels derived from the group activity of each cage are shown in Figure 6A. The waveform plot of the diurnal variation in activity over the two weeks is shown for each strain in Figure 6B, and average activity during day and night are shown in Figure 6C. Overall, we observe robust diurnal variation in activity in both males and females in all strains over a two week period that is highly consistent across the replicates. C57L/J and C57BL/6J have the highest level of activity during the day, while FVB/J has the lowest activity. Most strains, except C3H/HeJ and FVB/NJ, show robust differences between day and night activity, as expected. This is likely because C3H/HeJ and FVB/NJ carry the *Pde6b^rd1^* allele, which results in early onset retinal degeneration and is present in over 50 inbred mouse strains [98]. The *Pde6b* mutation is a model of human retinitis pigmentosa, a hereditary disease resulting in progressive photoreceptor loss, eventually leading to blindness [99]. All strains show an increase in activity at lights off, except Cast/EiJ which experiences an increase in activity several hours before lights turn off. This behavior in Cast/EiJ was described as early runner" and is a models of familial advanced phase sleep syndrome in humans, and mapped to mouse chromosome 18 [100, 101]. We performed statistical analysis of our data which showed significant effect of strain and light condition, but surprisingly no significant effect of sex (Figure S8A, pairwise *post-hoc* comparison in Figure S8B).

**Table 1:** Strains and animal used in the strain survey analysis. Groups of three mice were observed continuously in the DAX for two weeks.

|  | JR# | Age (weeks) | Animals (n) | Groups (n) |
| --- | --- | --- | --- | --- |
| **Female** |  |  |  |  |
| A/J | 000646 | 9.29 | 9 | 3 |
| BALB/cByJ | 001026 | 10.29 | 12 | 4 |
| BTBR T+ Itpr3tf/J | 002282 | 9.81 | 9 | 3 |
| C3H/HeJ | 000659 | 9.43 | 9 | 3 |
| C57BL/6J | 000664 | 9.14 | 9 | 3 |
| C57L/J | 000668 | 10.43 | 9 | 3 |
| Cast/EiJ | 000928 | 7.43 | 9 | 3 |
| DBA/2J | 000671 | 11.43 | 9 | 3 |
| FVB/NJ | 001800 | 11.43 | 9 | 3 |
| **Male** |  |  |  |  |
| A/J | 000646 | 8.38 | 9 | 3 |
| BALB/cByJ | 001026 | 8.43 | 9 | 3 |
| BTBR T+ Itpr3tf/J | 002282 | 9.14 | 12 | 4 |
| C3H/HeJ | 000659 | 12.29 | 12 | 4 |
| C57BL/6J | 000664 | 10.29 | 9 | 3 |
| C57L/J | 000668 | 10.43 | 9 | 3 |
| Cast/EiJ | 000928 | 8.43 | 9 | 3 |
| FVB/NJ | 001800 | 11.43 | 9 | 3 |

Next, we determined whether activity in the home cage correlates with activity in an open field. We reasoned that perhaps animals that show hypoactivity in the open field may show higher levels of activity in home cage environments. Alternatively, we hypothesized that activity levels in the open field correlate with home cage activity, i.e. certain strains are hypo- or hyperactive regardless of the environment. We compared home cage data with open field data with our previously published[37, 102]. The mean movement speed in the light and dark periods over a 14-day assay in the DAX home cages was compared to the mean distance traveled over a 55-min assay in the open field. We did not observe a significant correlation between activity in the open field and home cage activity in males or females during the day or night (Figure S8C). For instance, A/J is known to display very low activity in the open field in multiple data sets [32, 103, 104]. However, we find that in the home cage A/J have moderate activity similar to other strains (Figure 6C). We conclude that open field and home cage activity levels capture differing internal states in mice. We also conclude that the DIV Sys provides valuable insights into the animal behavior that is independent of commonly used tests such as the open-field.

#### 3.7.2 Unsupervised analysis of mHydraNet embedding

The DIV Data provides a stream of raw video data along with embeddings for individual frames in a sequence. We next sought to determine if the embedding in mHydraNet was able to distinguish various cage conditions (Figure 7A). Embedding vectors were extracted from the bottleneck of the mHydraNet for five clips from day and night time conditions with black and albino mice. To understand the representations, we clustered them along a dimensionally reduced plot. We can deduce a rough understanding of the clusters. Daytime frames with mice sleeping in a pile (Figure 7A, blue), daytime frames with active mice (Figure 7A, purple), nighttime frames with active mice (Figure 7A, red), nighttime frames with sleeping mice (Figure 7A, green), and lastly daytime frames in red light conditions (Figure 7A, orange). Such representations of image data processed with mHydraNet can serve many downstream tasks such as dataset curation, behavior analytics, among others.

#### 3.7.3 Zero-shot segmentation with SAM and Grounding DINO

Finally, we used DAX videos to create custom training data using zero-shot segmentation. Such an application can provide significant utility for experimental paradigms in novel environments or unencountered animal models to which mHydraNet was not been previously exposed. These datasets may serve both to retrain mHydraNet and to facilitate autonomous downstream analyses. In order to accomplish this, we used grounding DINO [105] combined with Segment Anything [106] and DEVA to propagate segmentation masks across videos in an unsupervised manner [107]. This allows us to track each animal (Figure 7B, top row) in a video stream (Figure 7B, bottom row). We qualitatively accessed several video clips and found good performance which can serve as a good starting point for further training of models (Supplementary Video 5).

## 4 Discussion

With this and a companion publication [44], we introduce two complementary tools for the advanced quantification of rodent behavior: JABS, an open-field platform, and the DIV Sys, a robust and scalable home cage monitoring system. Home-cage monitoring offers distinct advantages in husbandry, behavior annotation, and drug development. However, its widespread adoption has been hindered by the complexity of implementing an end-to-end solution, which requires the development of specialized hardware and sensors, efficient data management and compute resources, and sophisticated machine learning models for behavior analysis, along with the resources for training, maintenance, and support of scientists. Furthermore, the use of custom apparatus for data acquisition complicates the sharing and reuse of models and data, and essentially requires each laboratory or organization to start from scratch. These challenges are major barriers to democratization.

DIV Sys is a modern end-to-end platform for rodent home-cage monitoring that addresses these challenges. DIV Sys was designed by a multidisciplinary team of hardware and software engineers, machine learning scientists, cloud computing specialists, web designers, IoT experts, and behaviorists. The DAX system includes hardware that complies with the AAALAS rodent housing standards. Dual cameras for video data capture allow high-resolution imaging in visible and NIR lighting for day and night recordings. Video data is stored and processed in the cloud for behavior quantification using mHydraNet, a model optimized for both accuracy and speed in tracking multiple animals. The system also includes an associated study design and results visualization tool suite (DIV App) that enables collaborative animal study design and analysis. Scalable advanced machine learning tools are available using the DIV Data cloud platform for customized machine learning applications. The DIV Sys is built on modern software design principles, which ensures traceability of all analyses and supports the continual updating and refinement of analysis and ML models. We used the DIV Sys across six sites to analyze over a century of video data (2 pebibyte(PiB)). The vignettes presented here, in which we carry out a large strain survey of home-cage behavior in eight inbred mouse strains, and conduct clustering and segmentation mask creation for novel cage conditions, underscore the power and utility of this system.

### Potential stakeholders: animal care staff, drug developers, academics

The development of a digitally enabled platform for computer vision-based behavioral monitoring in laboratory animals has multiple applications across laboratory research areas including veterinary care, animal husbandry, pharma/biotech, neuroscientists, behaviorists, geneticists, pharmacologists, AI/ML practitioners, and data scientists.

The video technology within DIV Sys is an invaluable tool for the ethical management of laboratory animals. The system enables animal care staff to tag event notes and observations, and facilitates communication between animal technicians and veterinarians. Through the DIV App, staff can review the current and historical states of animals without the need for invasive manual health checks. Changes in activity levels can serve as indicators of animal health, and future developments will include automated health check metrics. For example, we have already developed a frailty scale using the JABS system. Many features from the visual frailty index could be adapted for use in DIV Sys to assess health status. Additionally, users can develop tools to detect adverse events such as seizures or cage flooding. As the user base expands and the system is trained on diverse datasets and conditions, these tools will continue to be refined.

In the drug development applications, the key advantage of DIV Sys is scalability with automated and objective measures. Traditional methods often struggle to handle complex behavioral phenotyping studies at scale. In contrast, the DIV Sys can scal to accomodate multi-cage stacks with cloud-based data analysis, enabling researchers to conduct more extensive, detailed, and reproducible studies. Furthermore, the platform supports auditable data sharing and standardization, which are critical for regulatory compliance in the pharmaceutical industry, particularly with the FDA.

For academic neuroscientists, behaviorists, geneticists, and pharmacologists, the DIV Sys provides a platform for continuous development, where work from one laboratory can be leveraged and extended by others. One example is exemplified by the creation of a frailty index to quantify biological age using JABS, which required almost 600 aged animals to train and validate. A uniform digital ecosystem across laboratories would allow others to utilize the trained frailty index model without the need to retest large numbers of animals. All models are versioned and traceable for reproducibility. Additionally, video data can be repurposed to train new frailty index models or other health metrics. We envision DIV Sys as a collaborative environment for such work across laboratories and organizations and a driver of innovation.

Finally, standardized data readily available at scale is ideal for AI/ML and data science researchers and algorithm developers. The integrated data science workbench further enhances scalability by providing computational scientists and biostatisticians with a robust platform to streamline and automate workflows related to database management and machine learning analysis. This not only increases efficiency, but also facilitates collaboration among team members through built-in annotation and study-sharing features. We envision a new generation of tool developers who operate in the DIV Sys ecosystem.

One key to the widespread adoption and accessibility of advanced phenotyping is the adoption of common behavior apparatuses that generate data in a standardized manner. Both JABS and DIV Sys have trained models for tracking of genetically diverse mice, thereby alleviating a major barrier to entry. For behavior annotation, JABS has an active learning system (JABS-AL) and a behavior classifier sharing app (JABS-AI) that can be easily incorporated into the DIV Sys. These allow users across sites to share data and behavior annotation.

### Limits of home-cage monitoring

The field of home-cage monitoring, including DIV Sys, has inherent limitations. We optimize cage design and conditions for effective video data acquisition. This requirement may present challenges in certain experimental paradigms where visual monitoring of the animal is not feasible or ideal. Currently, DIV Sys offers a limited set of behavioral metrics, including activity levels, wheel, water, and food region occupancy. In the future, we plan to expand these metrics to include those developed for JABS, for specific behaviors such as gait, posture, and grooming, and indices like frailty and nociception [33–35, 37]. Incorporation of novel sensors along with multimodal recording of behavior with neural activity and physiology would be valuable.

### Rodent homes and hotels: A model for community development

While home cage monitoring is effective in observing emotional, social, and homeostatic behaviors, it will not adequately capture all domains of neural function, such as cognitive or complex motor behaviors. Future extensions of the DIV Sys and JABS could include tasks designed to assess specific domains of behavior in an ethological manner and with additional sensors. Larger arenas such as the JABS open field arena can be modified to include these specialized tasks. For instance, JABS is large enough to test navigation tasks in a maze [108] or social tasks such as the visible burrow [109]. Animals could also be instrumented for neural recordings. A paradigm in which mice are continuously monitored in a home cage and periodically tested for specific domain function in a specialized arena (hotel) offers an attractive approach to link altered nervous system function and complex behavior over time. Indeed, mapping changes in behavior with altered neural and genetic circuits is a major task for the computational ethology community.

As a genetics institution, at The Jackson Laboratory, we see striking parallels between the field of advanced behavior quantification and the evolution of next-generation sequencing (NGS). Currently, advanced behavior quantification is at a stage similar to where NGS was in the mid-2000s. During that period, several NGS platforms were available but primarily confined to niche laboratories, making the technology largely inaccessible to the broader scientific community. It was not until the widespread adoption of the Illumina platform that NGS became more universally accessible. NGS is essentially a visual task performed on a microscopy platform. The details of image acquisition from solid-phase PCR and the subsequent conversion of images to sequences were managed by the platform itself. This allowed researchers to focus on leveraging the technology for innovation, such as developing novel sequencing libraries, sample preparation from various organisms and cells, and creating new data analysis approaches. The adoption of a common platform led to an explosion in sequencing innovation and to its current commodified use in laboratories and clinics.

Similarly, adoption of standardized behavioral analysis platforms has the potential to transform the field of behavioral research. Much like NGS, the approach allows modularization of this complex technology, which allows experts in various fields to work together while focusing on their area of strength. It enables biologists to focus on developing novel behavior assays that link altered behaviors to disease states, while relieving them of the burdensome and complex tasks of hardware creation, data handling, machine learning, and other back-end processes. Such computational tasks are critical and may be better suited for tool and algorithm developers. Data scientists and algorithm developers can work to apply their skills by accessing data at scale in order to develop a new suite of behavior tools. We recognize that this is a significant undertaking. To facilitate this transition, The Jackson Laboratory is committed to openly sharing the DIV Sys and model architecture, behavioral datasets, and statistical models with the research community.

## 5 Methods

### 5.1 Caging

We use a disposable rat cage (43.2 x 34.0 cm) with 909.7 cm^2^ floor space on the interior of the cage (InnoVive; San Diego, CA).

### 5.2 Environment measures

The assessment of the static environment was conducted at the Jackson Laboratory using two studies, each with five cages housing 3 Balb/cJ female mice each (20-24 weeks of age). Mice were housed on 50/50 blend of aspen chips (P.J. Murphy Forest Products Corp, Montveille, NJ) and aspen shavings Northeastern Products Corp, NEPCO, Warrensburg, NY) and had 1 cotton square nestlet (Ancare Corporation; Bellmore, NY) per cage. The power was shut down to the blower allowing the DAX cages to sit static. The first assessment was conducted for 7 hours and the second assessment was conducted for 8 hours. The goal was to run each assessment for 12 hours, however, due to equipment failure, the assessment periods were shortened. After 7 and 8 hours, the PortaSens II (Analytical Technology Inc,; Collegeville, PA) was used to assess carbon dioxide (CO_2_), oxygen (O_2_), ammonia (NH_3_), and the Vaisala M170 Measurement Indicator with an HM70 Humidity and Temperature Sensor (Vaisala Inc.; Vantee, Finland) was used to assess temperature and relative humidity within the cage (Figure S5). Another study was conducted at the Jackson Laboratory to assess the efficiency of the cage airflow component was tested over 6 weeks using 10 DAX cages housing 2 Balb/cJ female mice (6-8 weeks of age) per cage. Mice were housed on 50/50 blend of aspen chips (P.J. Murphy Forest Products Corp, Montveille, NJ) and aspen shavings Northeastern Products Corp, NEPCO, Warrensburg, NY) and had 1 cotton square nestlet (Ancare Corporation; Bellmore, NY) per cage. CO_2_, NH_3_, temperature, and relative humidity were measured weekly using the previously described sensor equipment over the course of 6 weeks (Figure S6).

### 5.3 Pathology

A study was conducted at The Jackson Laboratories (JAX) to examine the impact of LED lights at 100 LUX on rodent eye physiology. Twenty female BALB/cJ female mice (JAX) were housed 2 per cage in DAX cages for 10 weeks. Mice were humanely euthanized, eyes were collected, and the tissue samples were submitted for pathology evaluation. Pathological evaluations were conducted by a trained pathologist at JAX.

### 5.4 Strain Survey

All mice were ordered from the Jackson Laboratory production colony and immediately placed in the DAX upon arrival into our colony. Mice were housed on 50/50 blend of aspen chips (P.J. Murphy Forest Products Corp, Montveille, NJ) and aspen shavings Northeastern Products Corp, NEPCO, Warrensburg, NY) and had 1 cotton square nestlet (Ancare Corporation; Bellmore, NY) per cage. Mice were housed in groups of three mice per cage for the duration of testing. There was one death due to fighting behavior in male Cast/EiJ on the first day of recording. All animals were between 7.43 and 12.29 weeks at the start of the recording. Data from the time mice were placed into the DAX to the next onset of lights-on were treated as an acclimatization period and removed from analysis. Data was obtained from the DIV App as cage activity means in 1 hour bins that was normalized for the number of animals. Mean differences in animal activity were quantified using a one-way ANOVA test and a *post-hoc* Tukey test comparison. All analyses were performed using DIV Data tools.

### 5.5 Manufacturing of hardware

The hardware was designed and manufactured using industry standard design and development processes. Electronics were designed using the Altium suite of PCB design and layout tools. Mechanical components were designed using SolidWorks. A complete assembly of the system, including an export of the PCB from Altium, the cameras with virtual structures representing the field of view of the cameras, and all other sub-assembles provided confidence that the manufactured devices would meet all major requirements.

Once key parts had been prototyped for fit and functionality and the design had been reviewed by a team of engineers, a decision was made to produce 100 complete DAX devices. We engaged a contract manufacturer (http://aqs-inc.com) to produce a turnkey build of the devices. They acquired all of the necessary components and engaged subcontractors for key components, such as the sheet metal housing, display sub-assembly, and camera sub-assembly. Several components, such as the camera shroud and the fan mounts, were 3D printed at volume (http://re3D.org Gigabot) to avoid the cost of tooling molds. The contract manufacturer managed the entire supply chain and assembled the complete DAX. Our engineers worked closely with the contract manufacturer to ensure quality, focus the cameras and align the components, program the embedded software into the platform, and finally configure and test the complete system before shipping them to end users.

#### Hardware Parts

| Component | Manufacturer | Part Number |
| --- | --- | --- |
| Blower, Flow Sensor | Sensiron | SFM3013-300-CL |
| Blower, Fan | DELTA | BFB0512VHD-F00 |
| Blower, HEPA | CAMFIL USA, Inc. | P7-06.00-06.00-1-01-2B-00-0-0 |
| Visible LED, 5000K | Broadcom | ASMT-UWB1-NX3D2 |
| NIR LED, 940nm | Broadcom | AREQ-9020-MP000 |
| Light Sensor | Vishay | VEML7700-TR |
| Image Sensor | Sony | IMX477 / IMX577 |
| Center Camera Lens | Commonlands | CIL027 |
| Side Camera Lens | Commonlands | CIL068 |

## 6 Data and Code Availability

All relevant data and code will be available at https://github.com/KumarLabJax

## 7 Acknowledgments

We thank members of the Kumar Lab for critical feedback, including Brian Geuther, Gautam Sabnis, Jaycee Choi, Anshul Choudhary, and Jacob Bieirle. We thank Dr. David Coleman (JAX) for mouse pathological evaluations. We thank Marcin Klapczynski for figure design. We thank Victor Gehman for input on various aspects of the DIV Sys. We thank Anna Lisa Lucido for critical manuscript design feedback.

## 10 Supplement

**Figure S1:**
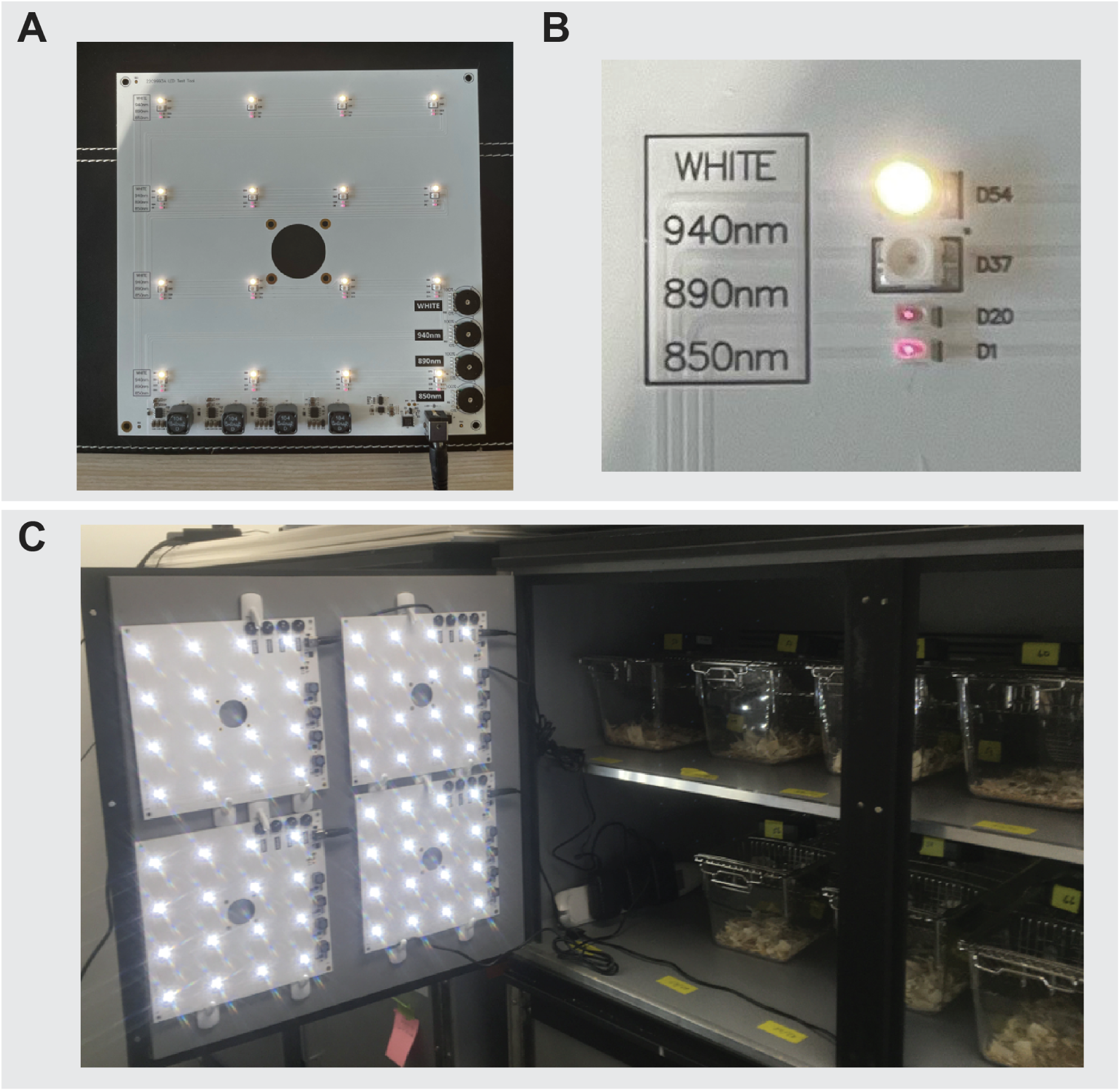
Light Test Design. (A) An image of the entire light board with three wavelengths of IR light (850nm, 890nm, 940nm) as well as white light. There are 16 light unites on each board with adjustable power knobs (bottom right) (B) Image of an individual light unit on the board. All lights are set at 100% power. The white light, 850nm, 890nm are visible to the camera, while the 940 is invisible. (C) An image of the wheel running light cabinets with 4 light boards on each side. The light power was cycled (LD 12:12) using a timer.

**Figure S2:**
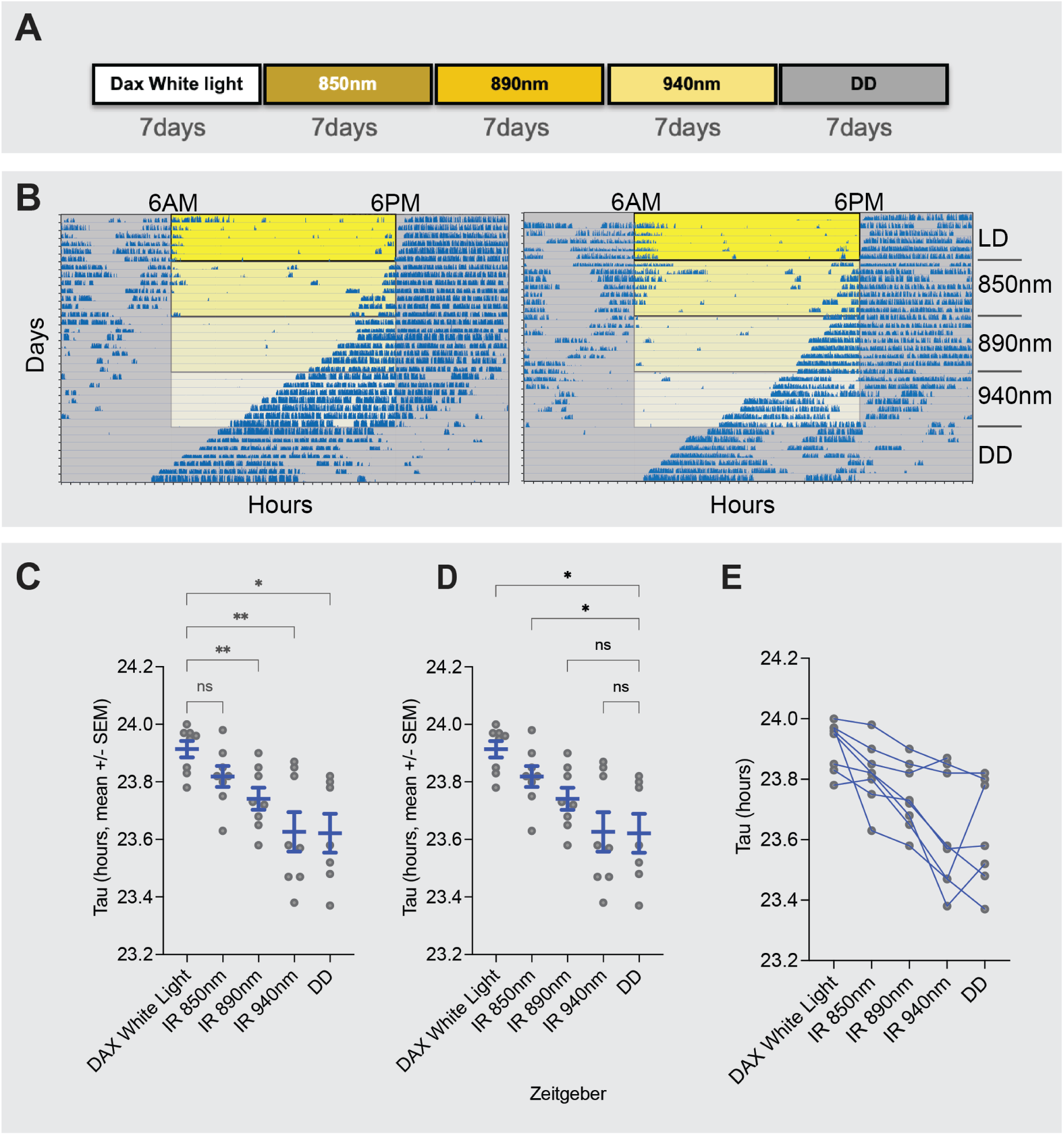
Masking of wheel running behavior by various infrared wavelengths of light. 850nm and 890nm have clear masking of activity, while 940nm IR light has no masking effect. Wheel running experiment was designed as shown in Figure S1(A) The light schedule. (B) Sample actograms of two animals. The colors indicate altering light wavelengths. (C-E) Quantification of period (Tau) for each animal at varying wavelengths. The same data are shown in all three plots. Mixed-effects model for repeated measure, Dunnetts test to control for multiple comparisons. (A) uses LD Tau as control, (B) uses DD Tau as control, and (C) each animal is connected over light cycles.

**Figure S3:**
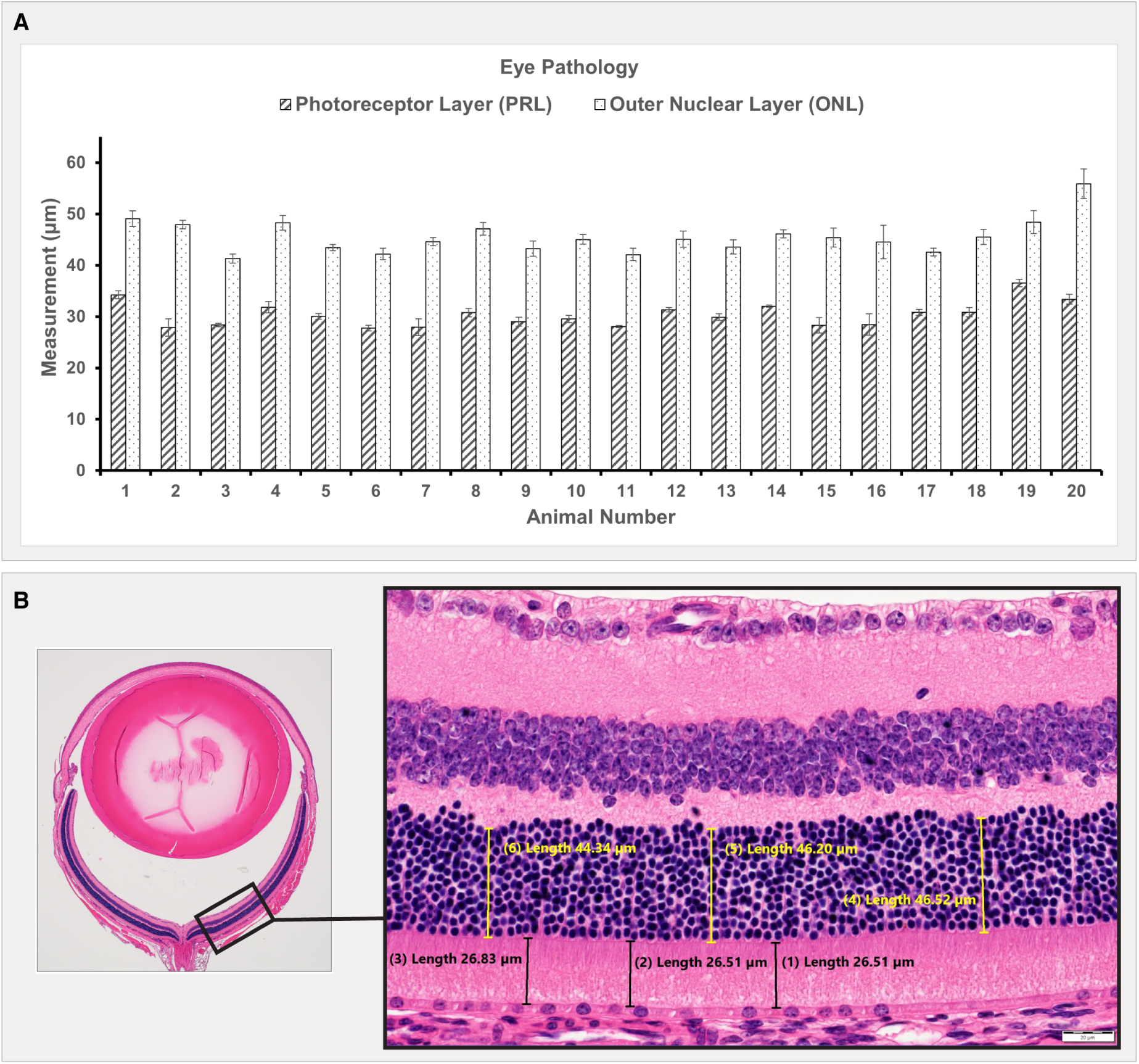
Histopathology analysis of retinas. (A) Thickness measurements of the photoreceptor layer and outer nuclear layer for each mouse. There is no evidence of retinal atrophy or other degenerative changes. (B) Measurements of the photoreceptor layer and outer nuclear layer, which contains the nuclei of photoreceptor cells. Several mice had eyes with developmental anomalies independent of lighting. There was no histologic morphology in the eyes of mice exposed to LED lighting. There is no evidence of retinal atrophy or other degenerative changes.

**Figure S4:**
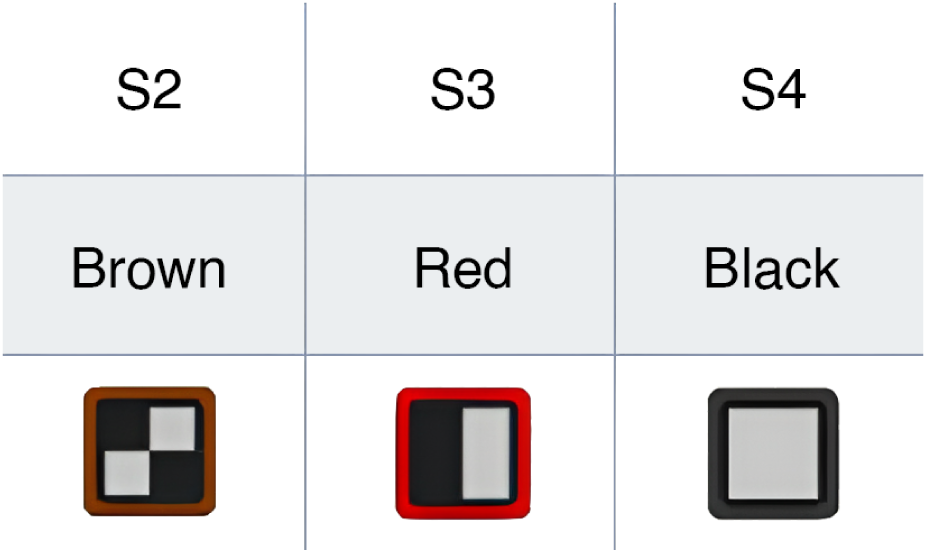
Ear Tags. Custom RapID Tagső (5mm) with 3 distinct 2-D matrix bar codes and colors were manufactured by RapID Lab, In (San Francisco, CA).

**Figure S5:**
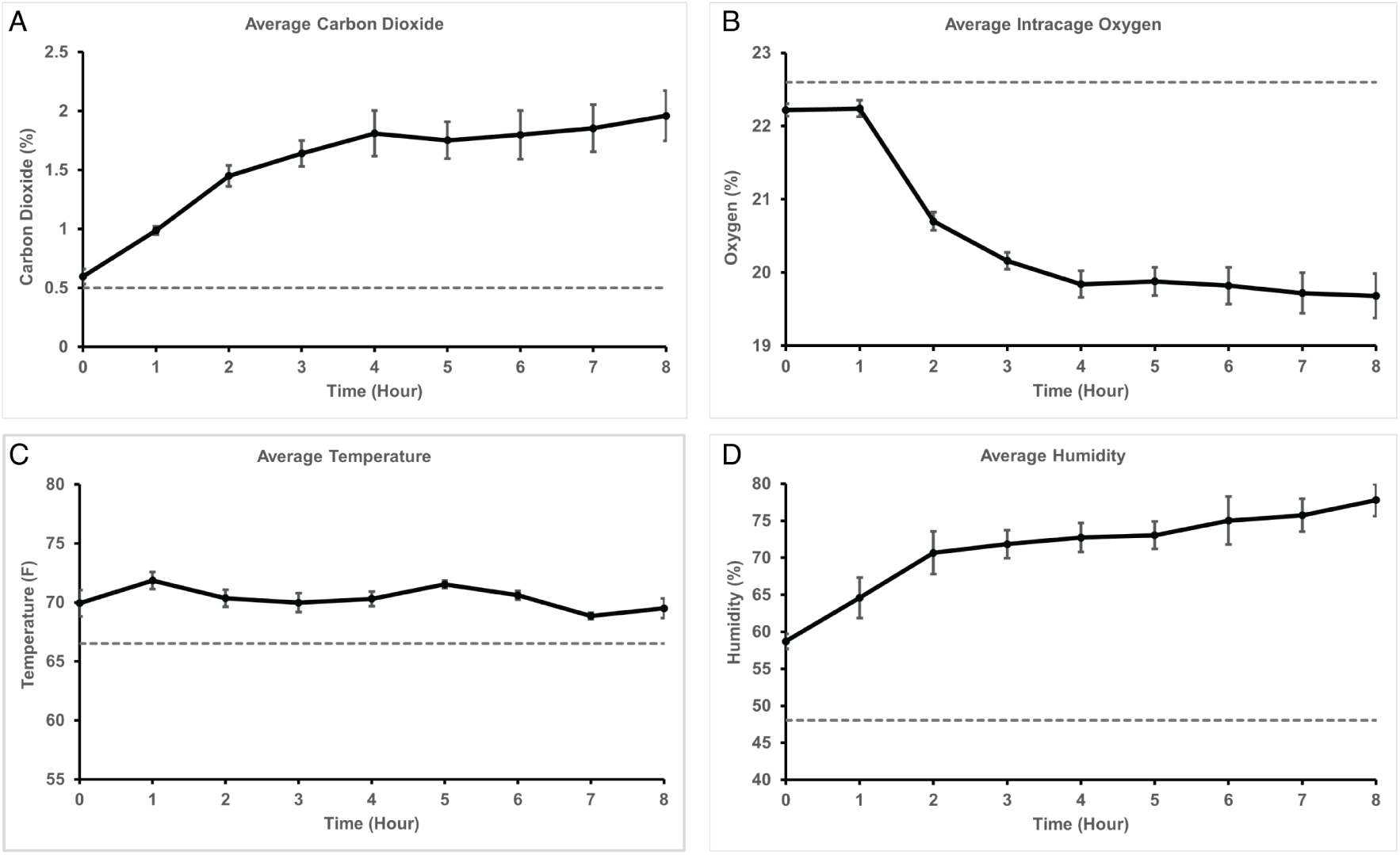
Static cage environmental testing by measuring environmental parameters without any power.(A) Carbon Dioxide (ppm), (B) Oxygen (percent), (C) Temperature (F), and (D) Humidity (percent) by time. Dashed lines indicate room reading. N = 10 cages. Error bars are SD.

**Figure S6:**
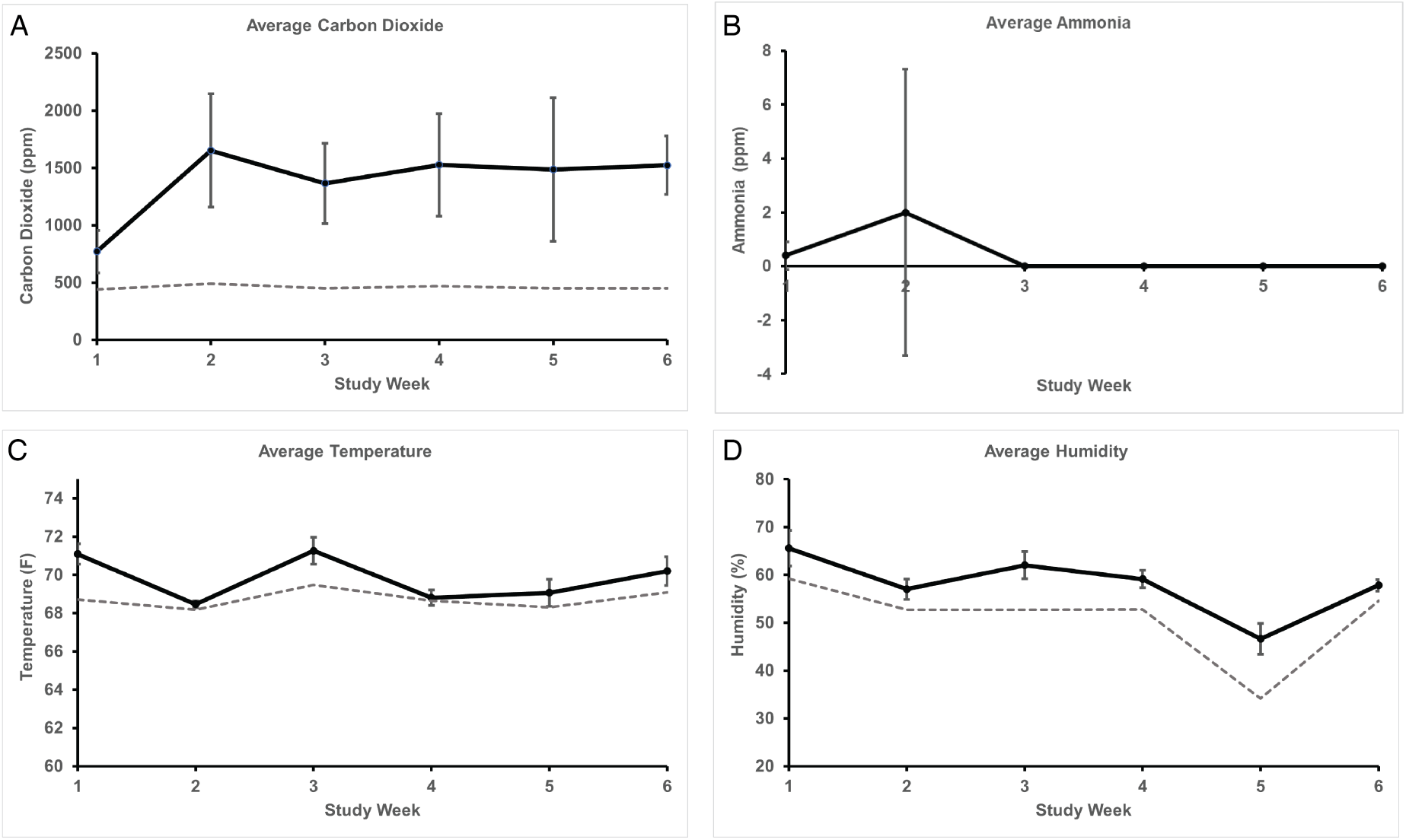
Long term cage study environmental testing. Carbon Dioxide (ppm), (B) Ammonia (ppm), (C) Temperature (F), and (D) Humidity (percent) by study week. Dashed lines indicate room reading. N=10 cages. Error bars are SD.

**Figure S7:**
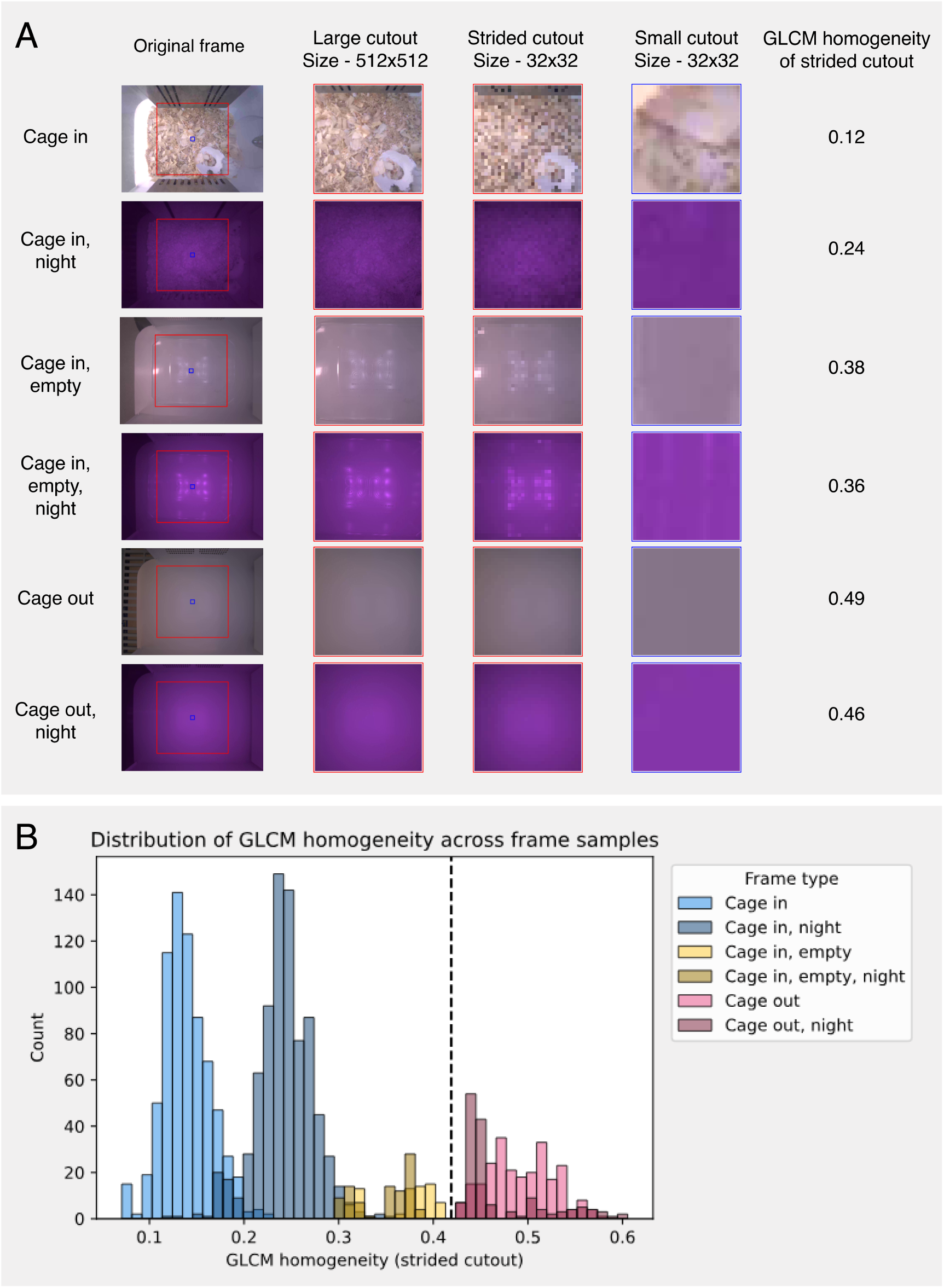
Cage in/out determination. (A) Example frames demonstrating different cage and lighting conditions. Red and blue squares are drawn on the original frame to illustrate the relative sizes of large (512 x 512 pixels) and small (32 x 32 pixels) cutouts. Strided cutouts, which sample every 16th pixel in both x and y dimensions, capture much of the textural information of a large cutout while requiring the reduced processing time of a small cutout. GLCM homogeneity values for these cutouts, shown to the right of each row, are greater when the cage is absent. (B) Histogram of GLCM homogeneity values for strided frame cutouts extracted from over 2000 frame samples. A homogeneity value of 0.41 allows for a separation of cage presence versus absence, with an area under the ROC curve of 1.0.

**Figure S8:**
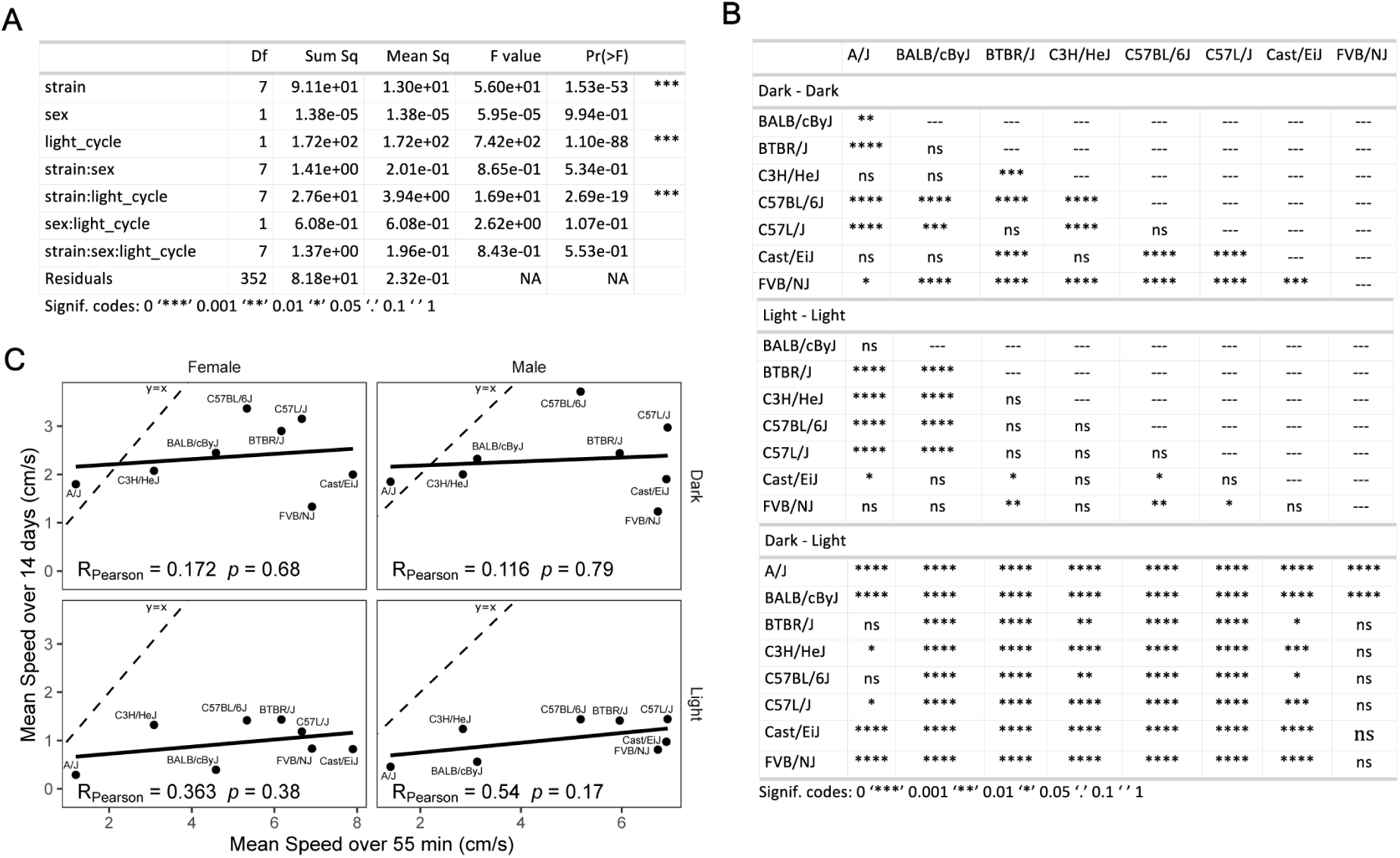
Strain survey statistics and correlation plots (A) One-way ANOVA table where average activity is characterized by strain, sex, light cycle, and the interaction between strain, sex, and light cycle. (B) A Tukey *post-hoc* comparison of inter- and intra-strain difference between light cycles. (C) Correlation scatter plots and Pearsons correlation coefficient for mean activity by strain, sex, and light cycle from the DIV-Sys compared to the mean activity represented in Kumar-MPD65002 [102]

